# Causes and consequences of linkage disequilibrium among transposable elements within eukaryotic genomes

**DOI:** 10.1101/2022.10.24.513493

**Authors:** Denis Roze

## Abstract

Sex and recombination can affect the dynamics of transposable elements (TEs) in various ways: while sex is expected to help TEs to spread within populations, the deleterious effect of ectopic recombination among transposons represents a possible source of purifying selection limiting their number. Furthermore, recombination may also increase the efficiency of selection against TEs by reducing selective interference among loci. In order to better understand the effects of recombination and reproductive systems on TE dynamics, this article provides analytical expressions for the linkage disequilibrium (LD) among TEs in a classical model in which TE number is stabilized by synergistic purifying selection. The results show that positive LD is predicted in infinite populations despite negative epistasis, due to the effect of the transposition process. Positive LD may substantial inflate the variance in the number of elements per genome in the case of partially selfing or partially clonal populations. Finite population size tends to generate negative LD (Hill-Robertson effect), the relative importance of this effect increasing with the degree of linkage among loci. The model is then extended in order to explore how TEs may affect selection for recombination. While positive LD generated by transposition generally disfavors recombination, the Hill-Robertson effect may represent a non-negligible source of indirect selection for recombination when TEs are abundant. However, the direct fitness cost imposed by ectopic recombination among elements generally drives the population towards low-recombination regimes, at which TEs cannot be maintained at a stable equilibrium.

## INTRODUCTION

Transposable elements (TEs) constitute an important fraction of the genetic material of many eukaryotes (Wells and Feschotte, 2020). Due to their capacity of self-replication, these genetic elements can indeed proliferate within genomes without bringing any net benefit to their host, earning them the qualification of “selfish DNA” (Doolittle and Sapienza, 1980; Orgel and Crick, 1980). While the number of copies of transposable elements per genome can reach very large values (for example, millions of copies of the *Alu* short interspersed element are present in the human genome, e.g., Cordaux and Batzer, 2009), their propagation is thought to be limited by two main factors: purifying selection acting against TE insertions, and the evolution of mechanisms restricting TE activity, which have been increasingly well described over recent years (Goodier, 2016; Kelleher et al., 2020; Almeida et al., 2022). Natural selection against TEs may take several forms. First, TE insertions may be deleterious when they occur within coding or regulatory sequences: a number of human diseases are indeed due to such insertions impairing gene function (Hancks and Kazazian, 2016; Payer and Burns, 2019). A second potential source of selection against TEs stems from fitness costs resulting from their activity (Nuzhdin, 1999). Third, the presence of similar sequences at different genomic locations may cause ectopic recombination events that often lead to strongly deleterious genomic rearrangements (Montgomery et al., 1987; Langley et al., 1988; Hedges and Deininger, 2007). Ectopic recombination among TE insertions has indeed been observed in *Drosophila melanogaster* (Montgomery et al., 1991) and is also a cause of human genetic disease (Payer and Burns, 2019). Although the role of ectopic recombination in the containment of TEs has led to some debate (e.g., Bíemont et al., 1997; Charlesworth et al., 1997), several lines of evidence suggest that it may be an important factor: in particular, the frequency of individual TE insertions correlates negatively with their length (longer insertions are expected to be more prone to ectopic exchange) and with the number of elements from the same family present in the genome, in both *D. melanogaster* and humans (Petrov et al., 2003; Song and Boissinot, 2007; Petrov et al., 2011). Furthermore, TEs tend to accumulate in genomic regions with reduced recombination (e.g., Table 1 of Kent et al., 2017), which is often interpreted as a consequence of lower rates of ectopic recombination in these regions.

**Table 1:** Parameters and variables of the model.

|  |  |
| --- | --- |
| $u$ | Transposition rate |
| $v$ | Excision rate |
| $\alpha$ | Fitness effect of a single TE insertion |
| $\beta$ | Coefficient of synergistic epistasis among TEs |
| $\gamma$ | Curvature of the fitness function (when fitness is given by equation 8) |
| $f$ | Number of TE families |
| $\sigma$ | Rate of sex |
| $s$ | Selfing rate |
| $o = 1 - s$ | Outcrossing rate |
| $N$ | Population size |
| $L$ | Number of possible insertion sites (assumed very large) |
| $r_{ij}$ | Recombination rate between insertion sites $i$ and $j$ |
| $R$ | Chromosome map length (in Morgans) |
| $\delta R$ | Effect of recombination modifier on chromosome map length |
| $\theta$ | Effect of chromosome map length on the rate of synergistic epistasis among TEs (direct cost of recombination generated by ectopic exchanges) |
| $c$ | Direct fitness cost of recombination (independent of TEs) |
| $W$ | Fitness of an organism |
| $p_i$ | Frequency of TEs at site $i$ in the population |
| $D_{ij}$ | Linkage disequilibrium between TEs at sites $i$ and $j$ |
| $n, \bar{n}$ | Number of TE copies in an individual, and mean number of TE copies per individual |
| $\text{Var}(n)$ | Variance in the number of TE copies per individual |
| $\rho$ | Effect of genetic associations among loci on the variance in TE number per individual ($\rho = 1$ in the absence of effect) |

Additional factors may generate a negative correlation between recombination rate and TE density, however (Kent et al., 2017). Gene density is often lower in genomic regions with reduced recombination, and the risk that a transposable element inserts into a coding sequence may thus be less important in these regions. Accordingly, TE density correlates more with gene density than with local recombination rate in the self-fertilizing species *Caenorhabditis elegans* and *Arabidopsis thaliana* (Duret et al., 2000; Wright et al., 2003), possibly due to the fact that ectopic recombination is less frequent in selfing than it outcrossing species, as it seems to occur mostly between heterozygous insertions (Montgomery et al., 1991). Furthermore, reduced recombination may evolve as a side effect of DNA methylation and chromatin modifications involved in the silencing of TEs, raising the possibility that local decreases in recombination rates may result from higher TE densities (Kent et al., 2017). Last, selection is less efficient in low recombination regions due to the Hill-Robertson effect (Hill and Robertson, 1966), which may cause an accumulation of TEs in these regions if TE insertions tend to be deleterious on average. This last hypothesis was explored by Dolgin and Charlesworth (2008) using a simulation model, showing that substantial increases in TE density are predicted only when recombination is very low.

It is interesting to note that contrasting views on the effect of recombination (or genetic exchange in general) on the dynamics of TEs can be found in the literature. On one hand, sexual reproduction is predicted to favor the spread of TEs, since in an asexual population (and in the absence of horizontal transfer) TEs stay confined into the lineage in which they first appeared (Hickey, 1982; Zeyl and Bell, 1996; Zeyl et al., 1996). In this scenario, sex and recombination tend to decrease the efficiency of selection against TEs by reducing the variance in the number of TEs per individual. On the other hand, the absence of recombination may lead to TE accumulation through Muller’s ratchet (Muller, 1964; Arkhipova and Meselson, 2005a; Dolgin and Charlesworth, 2008): here, recombination tends to increase the variance in TE number (and in fitness) among individuals, by breaking negative linkage disequilibria (LD) among TEs. In agreement with this dual role of sex and recombination, Dolgin and Charlesworth (2006) showed that a transition from a sexual to an asexual mode of reproduction may either lead to the proliferation of TEs or to their elimination, depending on parameter values (population size in particular). However, a more detailed understanding of the possible effects of recombination of TE dynamics requires the development of analytical models assessing the relative importance of the different possibles sources of LD among TEs.

A related, but somewhat less explored question concerns the effect of TEs on the evolution of recombination. Although the fitness cost generated by ectopic recombination among TEs should select for lower recombination rates (Kent et al., 2017; Brand et al., 2018), polymorphic TE insertions may also indirectly favor recombination. In particular, Charlesworth and Barton (1996) argued that non-zero rates of recombination may be favored in the presence of TEs, for two different reasons. First, ectopic recombination generates negative epistasis (on fitness) among TE copies (since fitness should decline faster than linearly with the number of copies present in the genome), and classical models have shown that negative epistasis among deleterious mutations favors the maintenance of non-zero recombination rates (Charlesworth, 1990; Barton, 1995). Second, increasing the rate of ectopic recombination leads to stronger selection against TEs, so that a modifier increasing recombination should eventually benefit from a lower load of TEs. However, the arguments of Charlesworth and Barton (1996) were mostly verbal, and a quantitative analysis of the relative importance of these different effects is still lacking. Additionally, the Hill-Robertson effect has been shown to be a potentially important source of selection for recombination in finite populations (Barton and Otto, 2005; Keightley and Otto, 2006; Roze, 2021), and the possible contribution of TEs to this process remains to be quantified.

This article presents approximations for the linkage disequilibrium between pairs of polymorphic TE insertions, in a classical model in which TEs are maintained at a stable equilibrium between transposition and epistatic selection (Charlesworth and Charlesworth, 1983; Langley et al., 1988; Charlesworth, 1991). The results show that, while negative epistasis tends to generate negative LD among elements, transposition generates positive LD. At equilibrium and in very large populations, the effect of transposition predominates, generating an excess variance in the number of TE copies per individual due to positive associations among TEs. This excess variance is substantial only for very low recombination rates, but may be important in the case of partially clonal or partially selfing populations with low rates of sex or outcrossing. In finite populations, the Hill-Robertson effect tends to generate negative LD among TE insertions, and this effect may be stronger than the deterministic effects of transposition and epistasis in regimes where drift is important, particularly among tightly linked loci. While the Hill-Robertson effect among TEs tends to favor recombination, the direct fitness cost of ectopic recombination generally predominates at high recombination rates, while TEs are often not maintained at a stable equilibrium when recombination is too low.

## METHODS

### Model of TE dynamics

The basic model of transposon dynamics considered here is similar to the one used in Charlesworth (1991) and Dolgin and Charlesworth (2006, 2008), representing a diploid population of constant size *N* with discrete generations (the notation used in the paper is summarized in Table 1). Individuals carry two copies of a linear chromosome, along which TEs may insert at random: each element generates a new copy with probability *u* per generation, the new copy inserting at a random location (drawn from a uniform distribution along the chromosome) either on the same or on the homologous chromosome. TEs may also be eliminated by excision, occurring at a rate *v* per element per generation. The fitness *W* of an individual is a decreasing function of the number of TEs in its genome (*n*), given by:

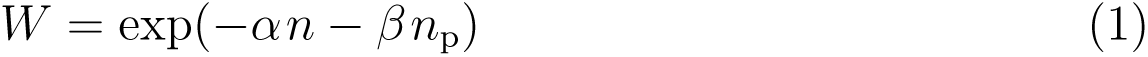

where *α* is the direct fitness effect of a transposon insertion, while the term in *β* corresponds to pairwise negative (*i.e.*, synergistic) epistasis among TEs (that may stem from the deleterious effect of ectopic recombination among elements), *n*_p_ being the number of pairs of TEs present at different sites in the genome. When *n* is large, and when each insertion at any given site stays rare in the population (and is thus mostly present in the heterozygous state), then *n*_p_ *≈ n*^2^*/*2 and equation 1 becomes equivalent to equation 1 in Dolgin and Charlesworth (2008). This model may also be seen as a particular case of Langley et al.’s (1988) model on the effect of ectopic recombination on TE dynamics, in the case where ectopic recombination is equally likely among all pairs of elements. Note that equation 1 does not take into account the fact that ectopic recombination may occur more frequently among heterozygous than among homozygous insertions (Montgomery et al., 1991). Introducing this feature into the model should not significantly affect the results when mating is random and insertions stay at low frequency, but would decrease the effect of selection against TEs under partial selfing, resulting in higher TE loads (Wright and Schoen, 1999; Morgan, 2001). The different events of the life cycle occur in the following order: excision – transposition – selection – reproduction (involving recombination and fertilization). During recombination, the number of crossovers occurring at meiosis is drawn from a Poisson distribution with parameter *R* (chromosome map length, in Morgans), the position of each crossover being drawn from a uniform distribution along the chromosome (no interference). Different reproductive systems will be considered: outcrossing with random mating, facultative sex (a proportion *σ* of offspring being produced by random mating, while a proportion 1 *− σ* is produced clonally), and partial self-fertilization (a proportion *s* of offspring being produced by selfing).

A modified model in which *f* different TE families are present in the population will also be considered. In this case, the rates of transposition and excision (*u* and *v*) are supposed to be identical for all families, while epistatic interactions (or ectopic recombination) only occur between elements from the same family, so that the fitness of an individual is given by:

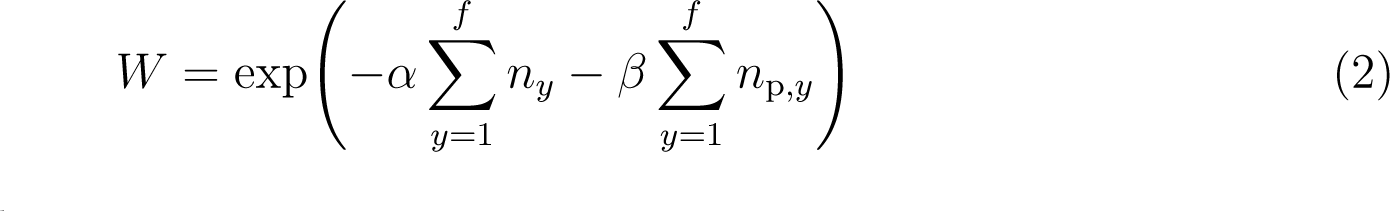

where *n_y_* is the number of TEs from family *y* in the genome of the individual, and *n*_p*,y*_ the number of pairs of TEs from family *y* present at different sites.

### Genetic associations and variance decomposition

Denoting *p_i_*the frequency of elements present at insertion site *i* in the population, the average number of elements per individual is given by *n* = 2 ∑*_i_ p_i_*. Furthermore, defining *X_i,_*_∅_ and *X*_∅_*_,i_* as indicator variables that equal 1 if an element is present at insertion site *i* on the first or second haplotype of an individual, and 0 otherwise, pairwise genetic associations can be defined as:

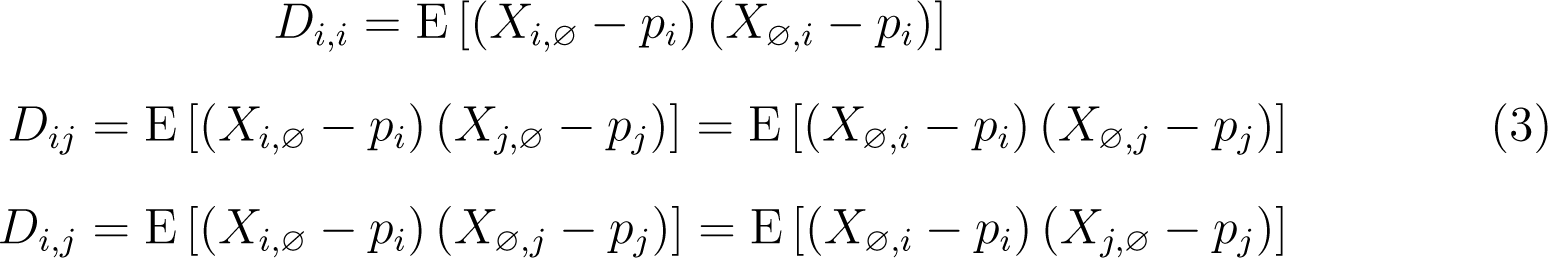

where E stands for the average over all individuals in the population (the last equalities of the last two lines of equation 3 stem from the fact that the model assumes no difference between sexes). The association *D_i,i_* measures excess homozygosity at site *i* (*D_i,i_* = 0 at Hardy-Weinberg equilibrium), while *D_ij_*corresponds to the classical linkage disequilibrium between sites *i* and *j* (on the same haplotype), and *D_i,j_*to the association between insertions at sites *i* and *j* present on different haplotypes. Using these variables, the variance in the number of elements per individual can be decomposed as:

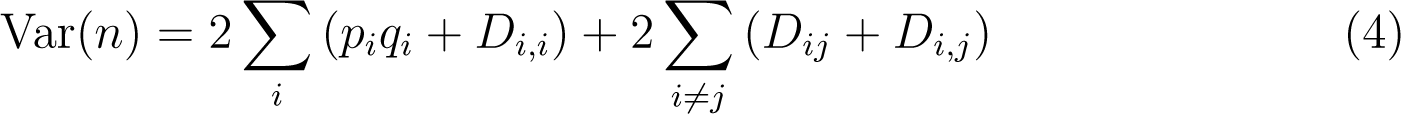

where the first sum is over all insertion sites, the second over all pairs of insertions sites, and where *q_i_* = 1 *− p_i_*. Assuming that the frequency of insertions is low at each site (so that *p_i_q_i_≈ p_i_*), and using the fact that *D_i,i_*= 0, *D_i,j_*= 0 under random mating, equation 4 simplifies to:

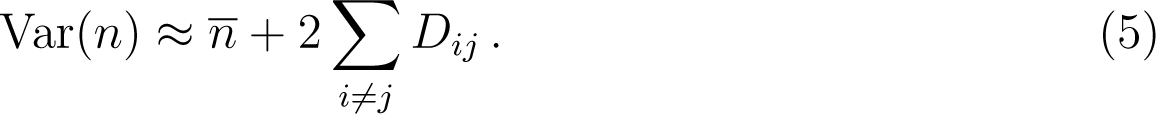

When linkage disequilibria (LD) are negligible, the number of TEs per genome is approximately Poisson distributed, with a variance equal to the mean (Charlesworth and Charlesworth, 1983). Positive LD tend to increase the variance in TE number (increasing the efficiency of selection against TEs), and negative LD to decrease the variance (decreasing the efficiency of selection). The overall effect of LD on the variance can thus be measured as:

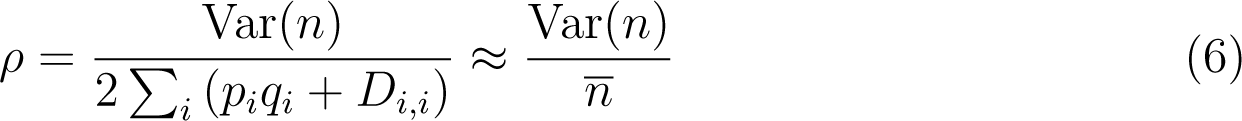

that equals 1 when the effect of LD is negligible. Under partial selfing, *D_i,i_* = *F p_i_q_i_* where *F* is the inbreeding coefficient (close to *s/* (2 *− s*) when selection is weak at each site) while associations *D_i,j_* may differ from zero. In that case, the effect of genetic associations between loci on the variance in TE number is given by:

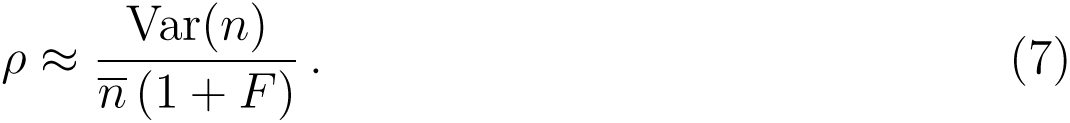

The same expression can be used under partial clonality, except that *F* equals zero in this case (at least when population size is sufficiently large, so that drift can be ignored). In the Supplementary Methods, approximations for *ρ* are derived under the different life cycles considered, assuming that the parameters *u*, *v*, *α* and *β* are small.

### Simulation programs

Individual based simulations are used to check the validity of analytical approximations. The simulation program (written in C++ and available from Zenodo) represents a population of *N* diploids, each carrying two copies of a linear chromosome (represented as an array of real numbers between 0 and 1, corresponding to the positions of TEs present on the chromosome). At the start of the simulation, the number of TEs in each individual is drawn from a Poisson distribution with parameter *n*_init_ (generally set to 10), their positions along the chromosome being drawn from a uniform distribution. The different events of the life cycle occur as in the analytical model. Three different outcomes are possible depending on parameter values: (i) TEs are eliminated from the population, (ii) the number of TEs increases indefinitely (in which case the program must be stopped), (iii) the number of TEs fluctuates around an equilibrium value (equilibrium being reached during the first 10^5^ generations for most parameter values used in the paper). In the last case, simulations generally run over 10^6^ generations, the mean and variance in the number of TEs per individual being measured every 100 generations (averages are computed over the last 9 *×* 10^5^ generations). The program also estimates the quantity ∑*_i_ p_i_*^2^ from the mean number of TE insertions shared between two chromosomes (over 1000 pairs of chromosomes randomly sampled from the population), so that the effect of genetic associations between sites on the variance in TE number can be computed as Var(*n*) */* (2∑*_i_ p_i_q_i_*) = Var(*n*) */* (*n −* 2∑*_i_ p_i_*^2^) under random mating and partial clonality, and Var(*n*) */* [(*n −* 2∑*_i_ p_i_*^2^) (1 + *F*)] under partial selfing (however, neglecting the terms in ∑*_i_ p_i_*^2^ often yields similar results). In a different version of the program, pairwise LD are measured over different classes of genetic distance between elements, these LD measures being taken every 1000 generations, from 100 replicate samples of 200 individuals. For a given set of parameters, the program must be run multiple times in order to obtain enough data points for tightly linked TEs. A third version of the program considers different shapes of fitness function: in this case, fitness is defined as:

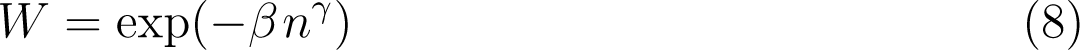

where *γ >* 1 to ensure that TEs can be maintained at a stable equilibrium. Similar forms of fitness function have been considered previously (e.g., Charlesworth and Charlesworth, 1983).

When *f* different TE families are segregating in the population and under random mating, the variance in the total number of elements present in an individual can be decomposed as:

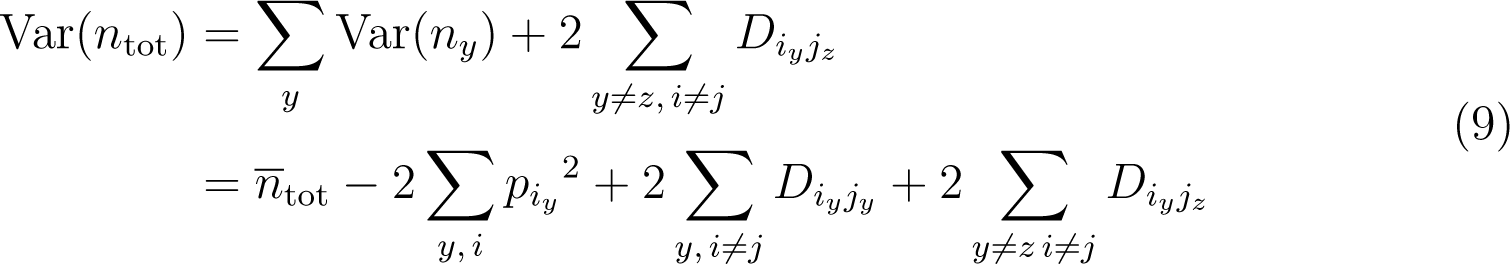

where *n̄*_tot_ and Var(*n̄*_tot_) are the mean and variance in the total number of TEs per individual, *n_y_* the number of elements from family *y*, *D_i__yjz_* the linkage disequilibrium between an element from family *y* present at site *i* and an element from family *z* present at site *j*, and *p_i__y_* the frequency of elements from family *y* at site *i*. The program thus measures 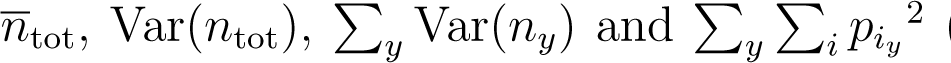 (estimated from the mean number of insertions shared between pairs of chromosomes) in order to infer the sum of LD between elements from the same family (second sum in the second line of equation 9), and the sum of LD between elements from different families (third sum in the second line of equation 9).

### Evolution of recombination

In the case of a randomly mating population, the effect of LD among TEs on selection for recombination is explored by incorporating a modifier locus affecting the genetic map length *R* of the chromosome. Two alleles *M* and *m* are segregating at this locus, the chromosome map length being *R*, *R* + *δR/*2 and *R* + *δR* in *MM* , *Mm* and *mm* individuals, respectively. In the deterministic limit (infinite population), the effect of linkage disequilibria between pairs of TEs on indirect selection at the modifier locus can be computed by extending the method of Barton (1995). In the case of a finite population, a general expression for the strength of selection for recombination generated by the Hill-Robertson effect between pairs of deleterious mutations (derived in Roze, 2021) can be transposed to the present model, as explained in the Supplementary Methods. These deterministic and stochastic terms can then be integrated over the chromosome to quantify the overall strength of indirect selection at the modifier locus (neglecting the effect of genetic associations involving more than 2 insertion sites). While ectopic recombination induces a fitness cost associated with recombination (generating a direct selective pressure to reduce *R*), this direct cost depends on the correlation between the rate of meiotic recombination and the rate of ectopic recombination. This correlation is controlled by a parameter *θ*, rewriting the epistasis coefficient *β* in equation 1 (that may be considered as a rate of ectopic recombination) as:

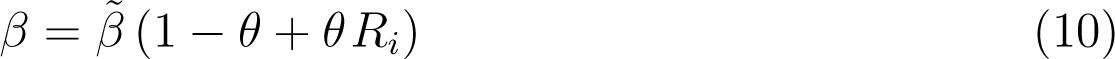

where *R_i_* is the chromosome map length of the individual (that depends on its genotype at the modifier locus and may take any positive value). When *θ* = 0, the rate of ectopic recombination is thus independent of *R* (no direct cost of recombination), while increasing *θ* increases the magnitude of direct selection against recombination due to the deleterious effect of ectopic exchanges among TEs. A direct fitness cost *c* par crossover (independent of ectopic recombination) may also be introduced into the model by multiplying the fitness of individuals by a factor *e^−cRi^* . Analytical predictions for the evolutionary stable chromosome map length are compared with individual based simulation results. For this, the simulation program is extended to include a modifier locus affecting *R*, using a similar setting as in Roze (2021) — see section 11 of Supplementary Methods for more details.

## RESULTS

### Deterministic model

In a randomly mating population, neglecting the effect of drift and linkage disequilibria, the average number of elements per individual at equilibrium is given by:

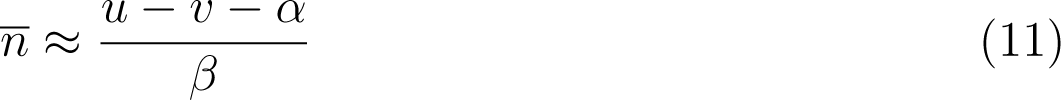

(Charlesworth, 1991, see also section 2 of Supplementary Methods), showing that negative epistasis (*β >* 0) is required in order to maintain the number of TE copies at a stable equilibrium. In section 3 of Supplementary Methods, the linkage disequilibrium between segregating elements at insertion sites *i* and *j* is shown to be approximately:

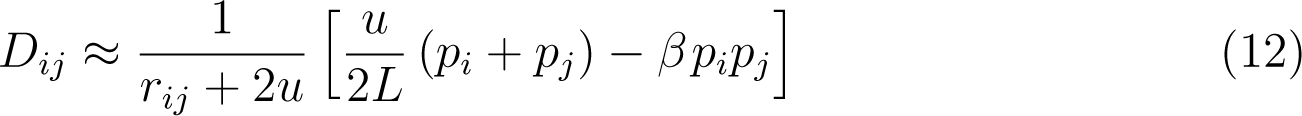

where *r_ij_*is the recombination rate between the two sites, and *L* the total number of possible insertion sites (assumed to be much larger than the number of elements present in a genome). Equation 12 shows that two different effects generate linkage disequilibrium among TEs: negative epistasis tends to generate negative LD (term in *β* in equation 12), while transposition generates positive LD (term in *u*). While the effect of epistasis on LD has been known for long (e.g., Felsenstein, 1965), the effect of transposition has not been formally described before. This effect stands from the fact that the insertion of a new TE copy into the same chromosome as the parental copy generates positive LD among the two copies. By contrast, when the new copy inserts into the homologous chromosome (say at site *j*), positive LD is generated if the individual carries the insertion at the initial site (say site *i*) in the homozygous state, while negative LD is generated if the individual is heterozygous for the insertion at site *i*. Because the individual is homozygous at site *i* with probability *p_i_* (and heterozygous with probability 1 *− p_i_*), the LD generated by transposition to the homologous chromosome is zero on average. This may be seen from the fact that *D_ij_* = *p_ij_ − p_i_ p_j_* where *p_ij_* is the frequency of chromosomes carrying insertions at sites *i* and *j*. New insertions at site *j* originating from copies at site *i* present on the homologous chromosome arise at rate *p_i_u/* (2*L*), thus increasing *p_j_* by this quantity, while *p_ij_* is increased by the fraction *p_i_* of these insertions falling onto chromosomes that are already carrying a copy at site *i*: *p_ij_*is thus increased by *p_i_ × p_i_u/* (2*L*), and the net effect on transposition to the homologous chromosome is zero. The overall effect of transposition is thus to generate positive LD among TEs, due to insertions into the same chromosome as the parental copy. One may notice that *u* also appears in the denominator of equation 12; this corresponds to the fact that directional selection against deleterious alleles tends to reduce LD by eliminating those alleles (e.g., Charlesworth, 1990; Roze, 2021), while TE excision also reduces LD. As shown in the Supplementary Methods (equations A29, A32), the strength of directional selection against TEs is *≈ u − v* at equilibrium, and LD is decreased by a term 2 (*u − v*) + 2*v* = 2*u* due to directional selection and excision.

Summing over all possible pairs of insertion sites and using *n̄* = 2 *_i_ p_i_*, equations 11 and 12 yield:

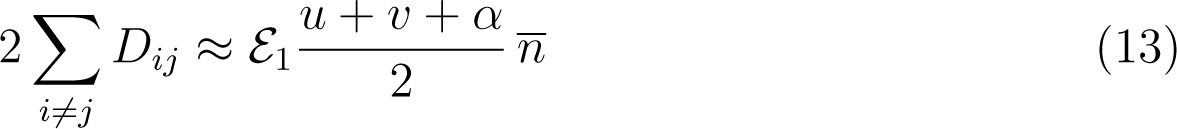

where *E*_1_ is the average of 1*/* (*r_ij_* + 2*u*) over all pairs of sites — an approximation for *E*_1_ as a function of *u* and *R* is given by equation A39 in the Supplementary Methods in the case where the density of crossovers is uniform along the chromosome. Equation 13 shows that in the present model, the effect of transposition (generating positive LD) is always stronger than the effect of epistasis (generating negative LD), so that LD is positive at equilibrium despite the fact that epistasis is negative. From equations 6 and 13, the inflation in the variance in number of TEs per individual due to positive LD can be expressed as *ρ* = 1 + *E*_1_ (*u* + *v* + *α*) */*2; however, it is shown in section 3 of Supplementary Methods that a more accurate approximation for the case of tight linkage (small chromosome map length *R*, so that *E*_1_ may be large) is given by:

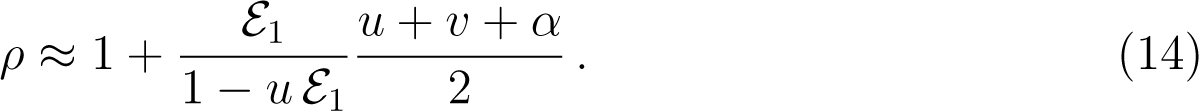

Figure 1 shows that equation 14 provides accurate predictions for the inflation in variance caused by LD among TEs under restricted recombination when *Nu* is sufficiently large and the mean number of elements per genome at equilibrium sufficiently small (e.g., *Nu* = 100, *n* = 10, top right figure), while important discrepancies appear under tight linkage, when *Nu* is smaller and/or *n* is larger. As we will see in the next subsection, these discrepancies are caused by negative LD generated by the Hill-Robertson effect in finite populations, that may lead to TE accumulation for parameter values leading to high *n* (bottom figures). Based on diffusion theory, the different parameters of the model should affect the results mostly through the *Nu*, *Nv*, *Nα*, *Nβ* and *Nr_ij_* products (e.g., Ewens, 2004; Dolgin and Charlesworth, 2006, 2008), and the deterministic and stochastic approximations derived here can indeed be expressed in terms of these quantities. When summed over all pairs of insertion sites, and assuming that *R* is not too large (roughly, *R ≤* 1) so that *E*_1_ can be approximated by equation A39 in the Supplementary Methods, the results only depend on *Nu*, *Nv*, *Nα*, *Nβ* and *NR*.

**Figure 1.**
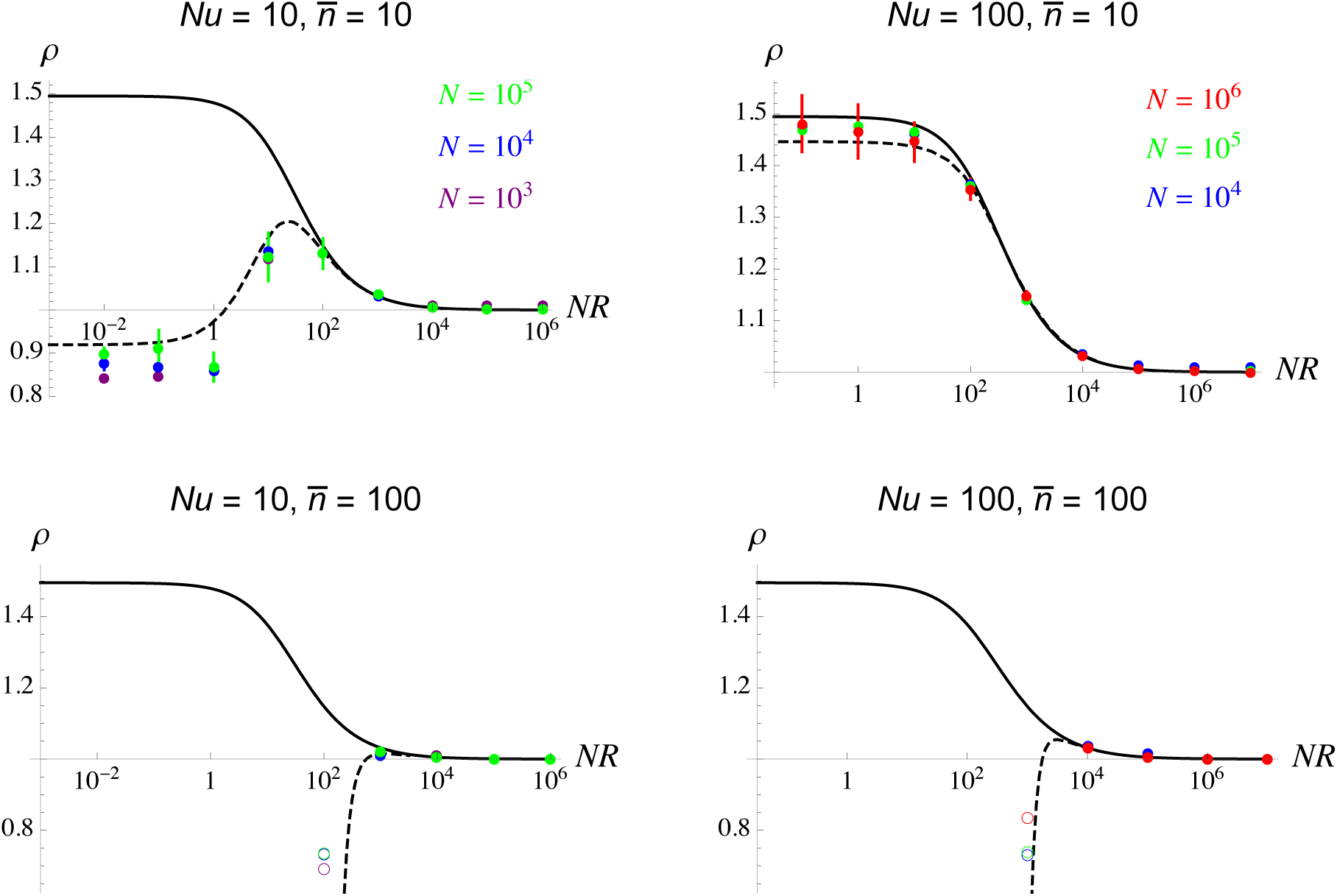
Effect of LD among TEs on the variance in TE number per individual (*ρ*, given by equation 6) as a function of the product of chromosome map length *R* and population size *N* (on a log scale), for different values of *Nu* and *n* (the mean number of TEs per individual at the deterministic equilibrium, given by equation 11). Solid curves: deterministic approximation (equation 14); dashed curves: stochastic approximation (including the Hill-Robertson effect, from equation A85 in the Supplementary Methods). Dots: simulation results; the different colors correspond to different values of population size *N* . Filled circles correspond to simulations during which the mean number of TEs per individual equilibrates; error bars (that are often smaller than the size of symbols) are obtained by dividing the last 9 *×* 10^6^ generations into 9 batches of 10^5^ generations and computing the variance over batch means. Empty circles in the bottom figures correspond to simulations during which TEs kept accumulating, and that had to be stopped (TE also accumulated in simulations with lower values of *R*); the circles correspond to averages over the last 10 data points of the simulations (last 1000 generations). Parameter values: *v* = *u/*100, *α* = 0, *β* = *u/*10 (top figures) or *u/*100 (bottom figures).

As can be seen in Figure 1, this is confirmed by the fact that simulations performed using different values of *N* (but keeping these products constant) lead to very similar outcomes. Note that a more accurate expression of *E*_1_ (valid for large *R*) can be obtained as explained in section 3 of Supplementary Methods, yielding better predictions for large (and rather unrealistic) values of *R*, as shown in Figure S1; however, *ρ* always stays close to 1 when *R* is large.

Intuitively, the effect of negative epistasis should become stronger when the curvature of the fitness function increases. Figure 2 shows the results of simulations in which fitness is given by equation 8, the parameter *γ* controlling the curvature. As can be seen on the figure, increasing *γ* (while adjusting *β* to ensure that *n* stays close to 10 under high recombination) indeed decreases the positive LD observed in the deterministic regime (high *Nu*), LD becoming negative when *γ* is sufficiently high. However, one can note that LD stays positive over a wide range of values of *γ*, including *γ* = 3. Figures 1 and 2 indicate that the effect of LD on the variance in TE number per individual becomes substantial only for values of *NR* that may seem unrealistically small for most populations, given that at least one crossover per bivalent occurs at meiosis in the majority of eukaryotic species (*R ≥* 0.5). However, even when *R* is high so that most pairs of insertion sites are at high genetic distance (with LD close to zero), LD between tightly linked pairs of TEs in the genome may be significantly positive in the deterministic regime (*Nu*, *n* small), as shown in Figure 3A for different values of *α* (the direct fitness cost of insertions). LD between tightly linked insertions becomes negative when the curvature of the fitness function is sufficiently strong, however, as shown by Figure 3B.

**Figure 2.**
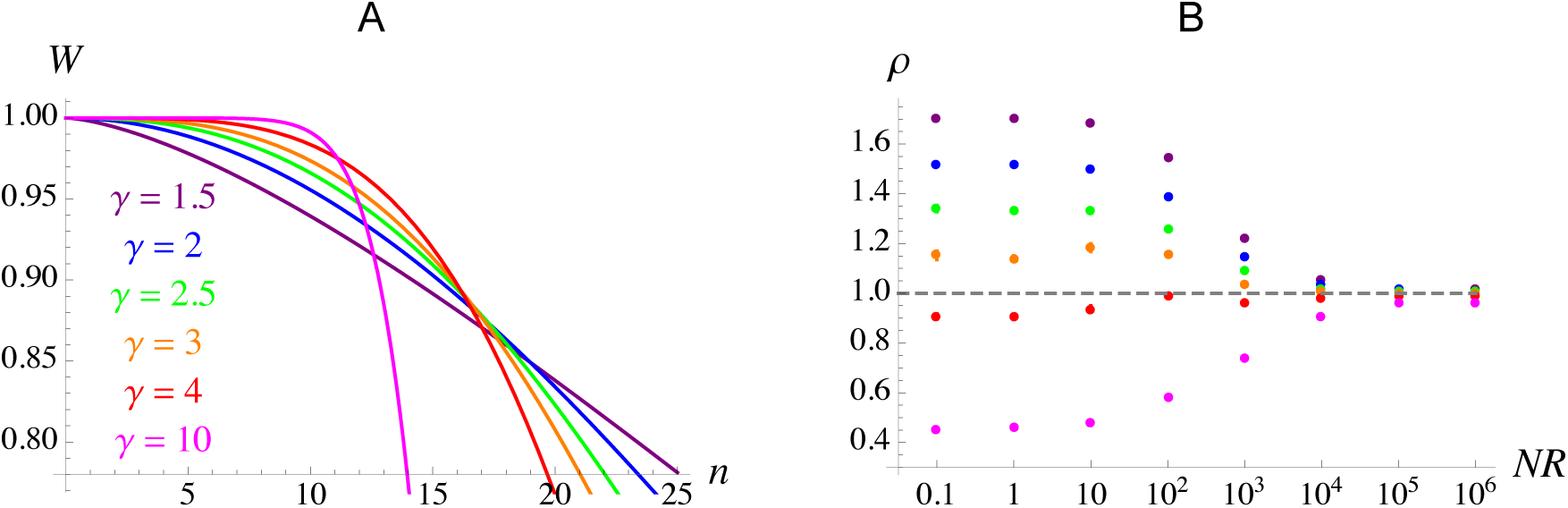
Shape of the fitness function (A) and effect of LD on the variance in TE number per individual as a function of *NR* (B) using the fitness function given by equation 8 and for different values of *γ*. Parameter values: *N* = 10^4^, *u* = 0.01, *v* = 10*^−^*^4^. For each value of *γ*, simulations were initially run with *R* = 10 and a range of values of *β* in order to determine the value of *β* leading to *n* = 10 by interpolation. This led to *β* = 1.98 *×* 10*^−^*^3^ (*γ* = 1.5), 4.54 *×* 10*^−^*^4^ (*γ* = 2), 1.09 *×* 10*^−^*^4^ (*γ* = 2.5), 2.67 *×* 10*^−^*^5^ (*γ* = 3), 1.65 *×* 10*^−^*^6^ (*γ* = 4) and 8.85 *×* 10*^−^*^14^ (*γ* = 10).

**Figure 3.**
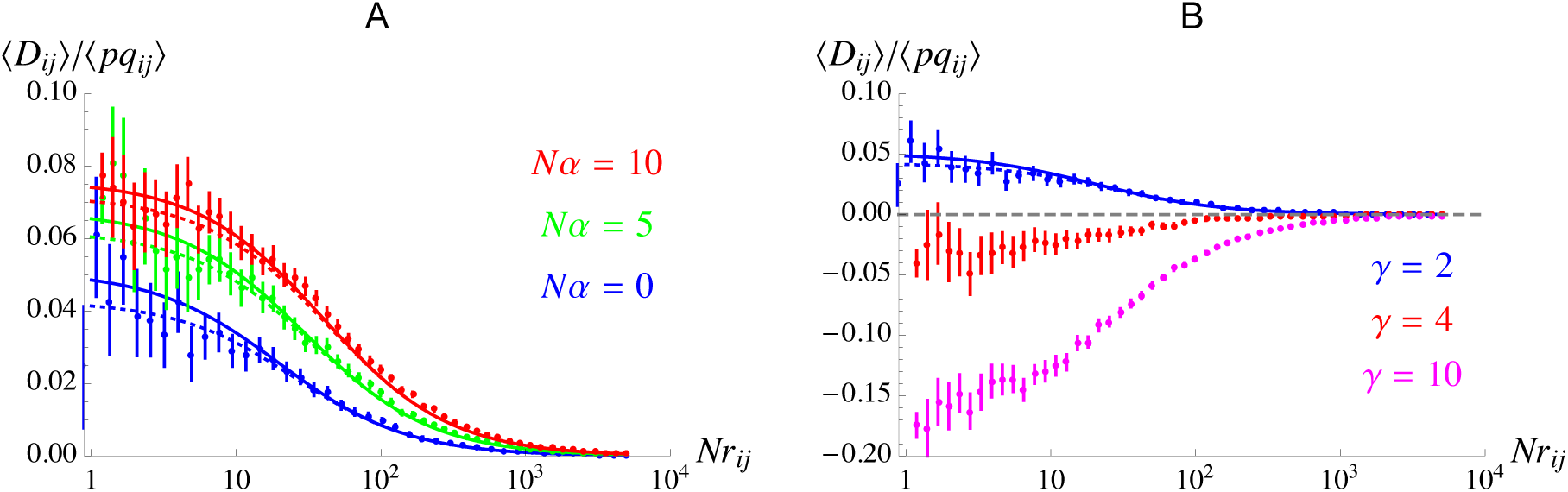
Average linkage disequilibrium between TE insertions at sites *i* and *j* divided by the average of *p_i_q_i_p_j_q_j_* (where *p_i_* is the frequency of insertions at site *i* and *q_i_* = 1 *− p_i_*) as a function of *Nr_ij_* (on a log scale). A: for different values of *Nα*, fitness being given by equation 1; B: for different values of *γ*, fitness being given by equation 8. Solid curves correspond to the deterministic predictions obtained from equation 12 (corresponding to the first term of equation 16), dashed curves to the stochastic approximation (including the Hill-Robertson effect) given by equation 16. In A, the transposition rate *u* is adjusted so that *n* = 10 according to equation 11, that is *Nu* = 10, 15, 20 for *Nα* = 0, 5, 10, respectively (with *Nβ* = 1). Dots correspond to simulation results for *N* = 10^4^ and *R* = 1. In the simulations, all pairs of segregating sites (in samples of 200 individuals taken from the population) are split into batches of log_10_*r_ij_*, *(D_ij_)* and *(pq_ij_)* being computed for each batch over 1000 points per simulation (one every 1000 generations, 100 replicate samples being taken for each point) and at least 100 replicate simulations. Error bars represent 95% confidence intervals obtained by bootstrapping over replicate simulations. In B, *u* = 0.001 and *β* is set to 1.71*×*10*^−^*^7^ and 6.84 *×* 10*^−^*^15^ for *γ* = 4 and 10 (respectively), so that *n ≈* 10 at equilibrium.

Positive LD generated by transposition may also substantially inflate the variance in number of elements per individual in facultatively sexual populations with a low rate of sex or in highly selfing populations. This is confirmed by the simulation results shown in Figures 4 and S2 for partial selfing, and in Figures S3 and S4 for partial clonality. As can be seen on these figures, contrarily to the deterministic approximation obtained under random mating (equation 14, that tends to a finite limit as the chromosome map length *R* tends to zero), the approximations obtained under partial selfing and partial clonality diverge (*i.e.*, tend to infinity) as the rate of sex or outcrossing decreases, and the inflation in variance caused by positive LD indeed reaches larger values in the simulations — up to nearly five-fold for *n* = 100 under partial selfing (Figures 4, S2), and ten-fold under partial clonality (Figure S4). However, the simulations also show that below a threshold rate of sex or outcrossing, the average number of TEs per individual cannot be maintained at a stable equilibrium anymore: TEs are either eliminated from the population (in deterministic regimes where the positive LD generated by transposition predominates) or accumulate indefinitely (in stochastic regimes where the negative LD generated by the Hill-Robertson effect takes over), TE accumulation occurring more frequently when the initial number of elements is larger and when *Nu* is lower (for a fixed *n*). This confirms the results obtained previously by Dolgin and Charlesworth (2006), showing that a transition from sexual to asexual reproduction may either lead to the accumulation of TEs or to their elimination depending on parameter values.

**Figure 4.**
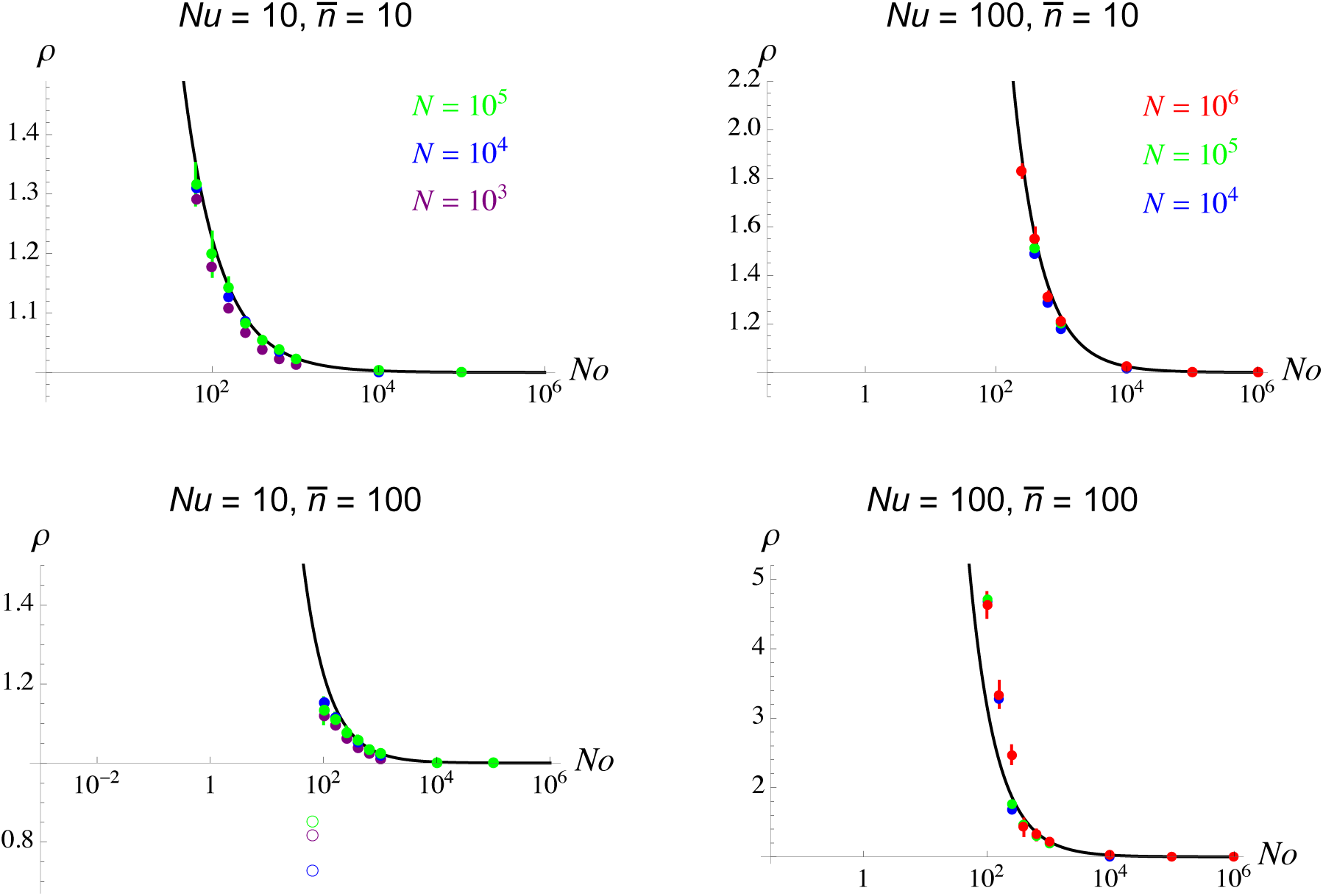
Effect of genetic associations among TEs at different sites on the variance in TE number per individual (*ρ*, given by equation 7) as a function of the product of outcrossing rate *o* = 1 *− s* and population size *N* (on a log scale) in a partially selfing population, for different values of *Nu* and *n* (the mean number of TEs per individual at the deterministic equilibrium under full outcrossing, given by equation 11). Curves correspond to the deterministic approximation given by equation A65 in the Supplementary Methods. Dots: simulation results; the different colors correspond to different values of population size *N* . On the left of the left-most filled circles of each figure, TEs are either eliminated from the population during the simulation or keep accumulating: TEs are eliminated in the top figures (*n* = 10), but accumulate in the bottom-left figure (*Nu* = 10, *n̄* = 100). As in Figure 1, the empty circles correspond to averages over the last 10 data points of the simulations. In the bottom-right figure (*Nu* = 100, *n* = 100) and when *No <* 10^2^, TEs are eliminated when the initial average number of elements per individual *n*_init_ is set to 10, but accumulate when *n*_init_ = 100. Parameter values: *R* = 10 (in order to mimic a genome with multiple chromosomes), *v* = *u/*100, *α* = 0, *β* = *u/*10 (top figures) or *u/*100 (bottom figures).

### The Hill-Robertson effect

As shown in section 2 of Supplementary Methods, the effective strength of selection against each TE copy is approximately *α* + *βn* under random mating, which (using equation 11) is *≈ u − v* at equilibrium (Charlesworth, 1991). When epistasis is weak relative to the effective strength of selection (*i.e.*, when *β « u − v*), previous results on the Hill-Robertson effect between deleterious alleles with independent fitness effects may be transposed to the present model. In particular, assuming that insertion frequencies *p_i_* stay close to their deterministic values, extending the analysis of Roze (2021) to the current model yields:

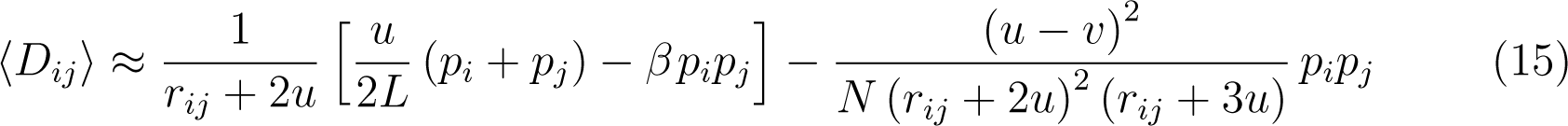

where *(D_ij_)* is the expected value of *D_ij_* at equilibrium (see section 6 of Supplementary Methods for more details). The second term of equation 15 corresponds to the Hill-Robertson effect (negative LD generated by selection and drift). Using *p_i_ ≈* (*u − v − α*) */* (2*Lβ*) (from equation 11) yields the following approximation for *(D_ij_) / (pq_ij_)*, where *(pq_ij_)* is the expected value of *p_i_q_i_p_j_q_j_*:

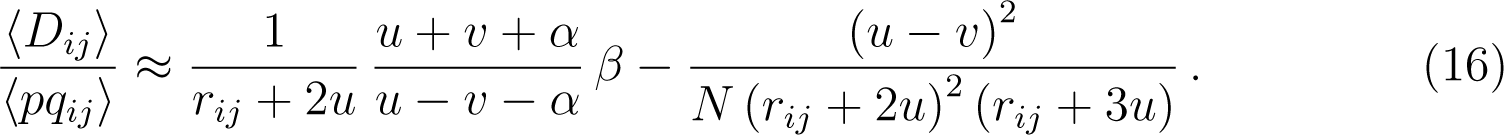

Note that this ratio does not depend on the number of possible insertion sites *L*. As shown by Figure 5, the relative importance of the Hill-Robertson effect increases as recombination decreases, as *Nu* decreases (for a fixed expected number of TEs per individual *n*) and as *n* increases (for a fixed *Nu*). While *(D_ij_) / (pq_ij_)* is often negative in the case of tightly linked loci, it may be positive when *Nu* is sufficiently large, for parameter values leading to moderate *n* (in this case, the deterministic effect of transposition predominates over the effects of epistasis and drift). Similar results are obtained in the case of partially selfing populations, LD between distant sites becoming more important when the selfing rate is high (Figure 5).

**Figure 5.**
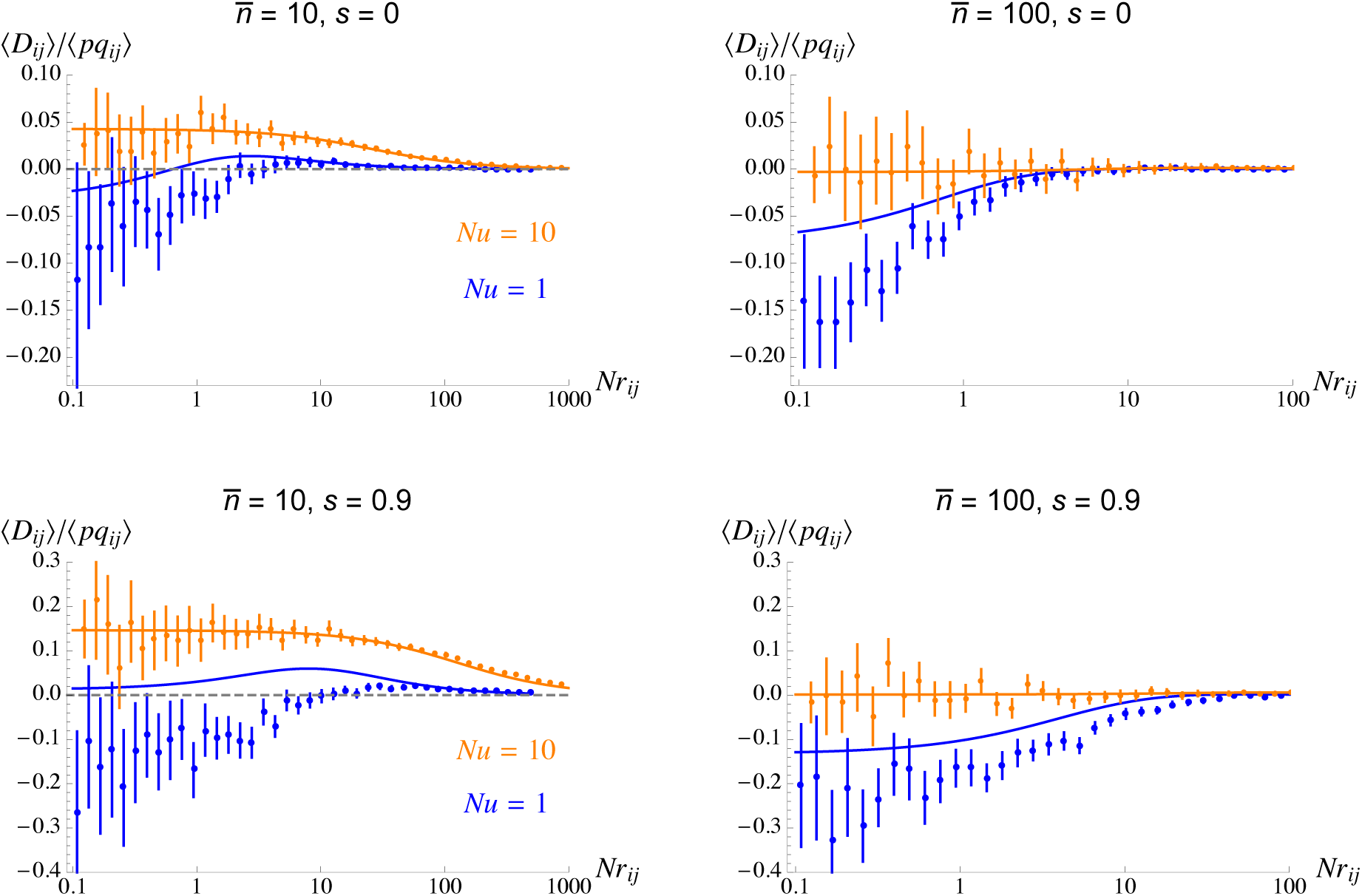
Average linkage disequilibrium between TE insertions at sites *i* and *j* divided by the average of *p_i_q_i_p_j_q_j_* as a function of *Nr_ij_* (on a log scale). Top figures: random mating; bottom figures: selfing rate *s* = 0.9. Curves correspond to analytical predictions from equations 16 (top) and A71 (bottom), and dots to simulation results for *N* = 10^4^ (orange) and *N* = 10^3^ (blue). In the simulations, all pairs of segregating sites (in samples of 200 individuals taken from the population) are split into batches of log_10_*r_ij_*, *(D_ij_)* and *(pq_ij_)* being computed for each batch over 1000 points per simulation (one every 1000 generations, 100 replicate samples being taken for each point) and large numbers of replicate simulations (up to 1000 for *N* = 10^3^). Parameter values: *v* = *u/*100, *α* = 0, *β* = *u/*10 (left) or *u/*100 (right), *NR* = 10^4^ (in the simulations).

Summing equation 15 over all possible pairs of sites yields the following approximation for the effect of LD on the variance in the number of elements per individual:

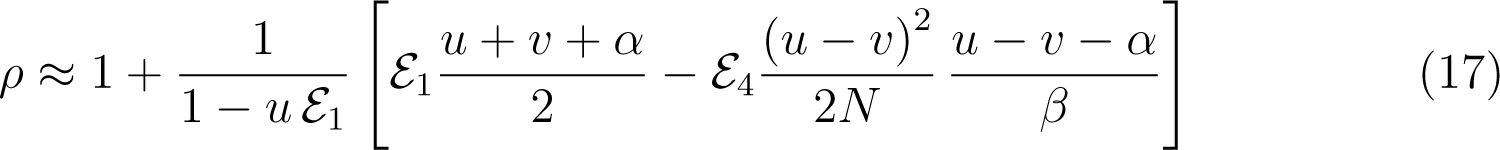

where *E*_4_ is the average of 1*/* (*r_ij_* + 2*u*)^2^ (*r_ij_* + 3*u*) over all possible pairs of insertion sites. Note that *N* in equation 17 should correspond to the effective population size *N*_e_, that may vary with respect to genomic location according to local interference effects among loci. With the genomic architecture considered here (single chromosome with map length *R* and a uniform density of crossovers), *N*_e_ may be significantly reduced by background selection effects caused by segregating TEs when *R* is small (Charlesworth, 1996), loci present in the central part of the chromosome being more strongly affected than loci located at the extremities. An analysis presented in section 7 of Supplementary Methods shows that the negative LD generated by the Hill-Robertson effect between TEs segregating at sites *i* and *j* is amplified by the presence of an element segregating at a third site *k*; the effect of background selection on *(D_ij_)* can then be approximated by integrating over all possible positions of site *k* along the chromosome. Using this method (that involves numerically integrating over all possible triplets of sites *i*, *j*, *k*) generates the dashed curves in Figure 1, that better match the simulation results than the deterministic approximation in the top figures (*n̄* = 10), and predict strongly negative LD in the parameter region where TEs accumulate in the bottom figures (*n* = 100 at the deterministic equilibrium).

The results shown above can be extended to the case where *f* different TE families are segregating in the population (see section 8 of Supplementary Methods). In this case, two forms of LD can be distinguished: between elements from the same family, or from different families. While the first type of LD (within family) is affected by transposition, epistasis and the Hill-Robertson effect (and may thus be positive or negative depending on the relative importance of deterministic and stochastic effects), the second type of LD (between families) is only generated by the Hill-Robertson effect and should therefore always be negative. This is confirmed by the simulation results shown in Figure 6, the approximations derived in the Supplementary Methods (equations A90–A92) providing correct predictions in regimes where the number of TEs per individual is maintained at a stable equilibrium. As can be seen on the bottom right figure, the sum of all linkage disequilibria (within and between families) is positive when *R* is sufficiently large (due to the positive LD within families generated by transposition), and becomes negative under weak recombination as the Hill-Robertson effect takes over. This transition occurs at higher values of *R* when the number of TE families *f* increases, due to stronger interference effects (the effective population size *N*_e_ decreases as *f* increases, since more elements are segregating). Note that the present model assumes no epistasis between TEs from different families; introducing negative epistasis between pairs of elements from different families would tend to make LD among them more negative. From the results shown in Figure 6, one expects that recombination should be disadvantageous when *R* is large (as the sum of all LD among TEs is positive and recombination thus decreases the variance in fitness), while recombination becomes advantageous at lower values of *R* (the sum of all LD becoming negative), in particular when the number of segregating elements is large.

**Figure 6.**
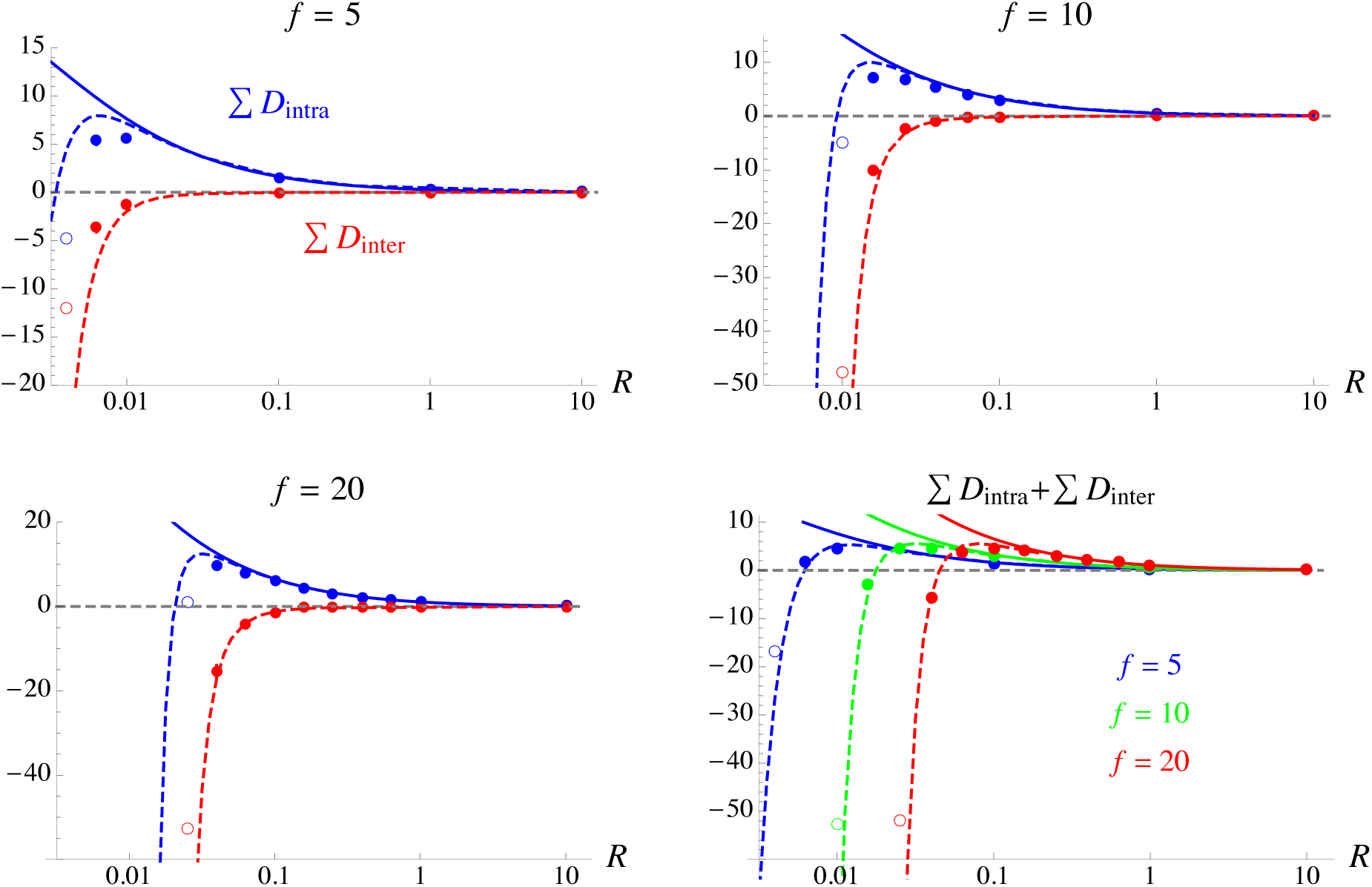
Top and bottom-left figures: sum of LD between pairs of elements from the same family (second sum in equation 9, blue), and sum of LD between elements from different families (third sum in equation 9, red), for different number of TE families segregating in the population (*f* = 5, 10, 20). Solid curves: deterministic approximation (equation A90 in the Supplementary Methods, without the term in 1*/N*); dashed curves: stochastic approximations (equations A90 and A92 in the Supplementary Methods); dots: simulation results (error bars are smaller than the size of symbols). As in Figure 1, empty circles correspond to simulations during which TEs kept accumulating (the circles corresponding to the average over the last 10 data points of the simulations). Bottom-right figure: sum of all LD among TEs (within and between families); curves and symbols have the same meaning as in the other figures. Parameter values: *N* = 10^5^, *u* = 10*^−^*^3^, *v* = 10*^−^*^5^, *α* = 0, *β* = 10*^−^*^4^ (*n* = 10 per family).

### Evolution of recombination

Two different types of selective force may act on a modifier locus affecting recombination rates: direct forces caused by direct fitness effects of the modifier, and indirect forces generated by linkage disequilibria between the modifier and selected loci. In the present model, a direct selective force against recombination is generated by the fitness cost caused by ectopic recombination among TEs, when meiotic recombination rates and rates of ectopic recombination are positively correlated (i.e., *θ >* 0 in equation 10). One may further assume a direct fitness cost *c* of crossovers (that may correspond for exemple to an energetic cost associated with the recombination process) independent of the presence of TEs. Indirect selection is generated by two different effects: (i) the disruption of LD among TEs by recombination, and (ii) stronger selection against TEs due to increased ectopic recombination, when *θ >* 0 (Charlesworth and Barton, 1996). These different selective forces are quantified in section 9 of Supplementary Methods, assuming that the chromosome map length *R* is not too small, so that the effect of LD among TEs on the variance in *n* among individuals remains weak (roughly, *NR >* 10^3^ for the parameter values used in Figure 1). This analysis shows that increasing the rate of ectopic recombination among TEs is always disadvantageous: although it allows a better purging of TEs (as well as other indirect effects described in section 9 of Supplementary Methods), this does not compensate for the direct fitness cost associated with ectopic recombination. Therefore, increasing the correlation between the rate of ectopic recombination and meiotic recombination rates (by increasing *θ*) disfavors recombination. By contrast, breaking LD among TEs may be beneficial or not depending on parameter values. In the deterministic regime where the Hill-Robertson effect is negligible, linkage dis-equilibria among TEs are positive, increasing the variance in the number of elements per genome. While breaking positive LD is beneficial in the short term when epistasis is negative (as it increases the mean fitness of offspring), it reduces the variance in fitness among offspring and thus decreases the efficiency of selection against TEs. From equations A100 and A104 in the Supplementary Methods, one obtains that the effect on the variance in fitness predominates over the effect on mean fitness in most cases, so that breaking positive LD is disfavored. The effect on mean fitness may be stronger than the effect on the variance when *R* is high and *n* is low (in which case breaking positive LD is beneficial), but indirect selection is then extremely weak and easily overwhelmed by any slight direct fitness effect of the modifier. By contrast, recombination is beneficial in regimes where the Hill-Robertson effect predominates, generating negative LD: in this case, recombination increases the variance in fitness and the efficiency of selection. Because LD tends to be positive when *R* is high and negative at lower values of *R* (e.g., Figure 6), indirect selection generated by LD among TEs is expected to favor intermediate recombination rates, at least when the number of TE copies per genome is not too small.

From the analysis presented in sections 9 and 10 of Supplementary Methods, the expected change in frequency of allele *m* (that increases the map length of the chromosome by an amount *δR*) can be decomposed as:

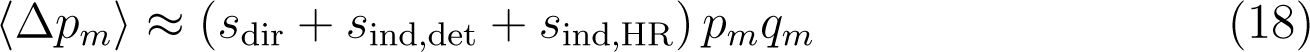

where *s*_dir_ is the strength of direct selection against recombination (due to the deleterious effect of ectopic recombination among TEs and to any inherent cost of recombination), and *s*_ind,det_, *s*_ind,HR_ the strength of indirect selection generated by deterministic and stochastic effects, respectively. Assuming that *δR* is small, *s*_dir_ is approximately:

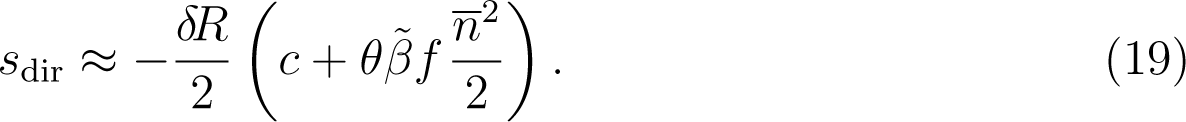

As discussed above, the deterministic component of indirect selection *s*_ind,det_ is often mostly driven by the negative effect of reducing the variance in fitness among offspring, in which case it can be approximated by (assuming *v*, *α « u*):

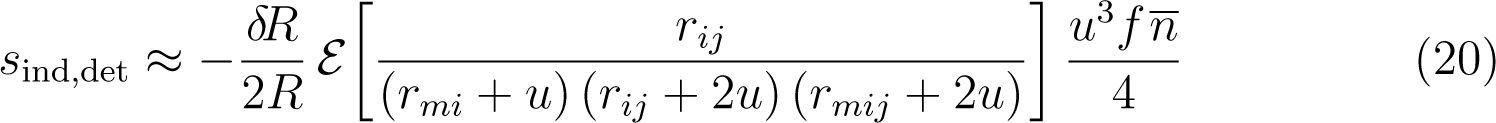

where *E* [*X*] is the average of *X* over all possible pairs of insertions sites *i* and *j*, and where *r_mij_* = (*r_mi_* + *r_mj_* + *r_ij_*) */*2 is the probability that at least one recombination event occurs among the three loci.

An approximation for the stochastic component of indirect selection *s*_ind,HR_ can be obtained from the expression derived in Roze (2021) for the strength of selection for recombination generated by the Hill-Robertson effect between deleterious mutations (see section 10 of Supplementary Methods). When *R » u*, one obtains:

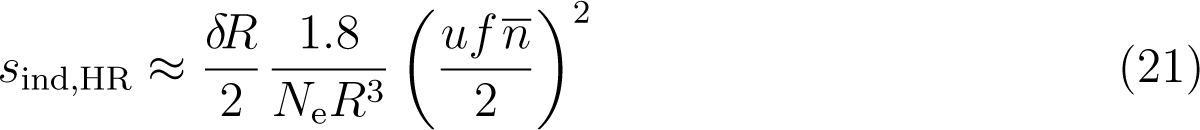

corresponding to equation 1 in Roze (2021), adapted to the present model. In-deed, the strength of selection for recombination generated by interference among deleterious mutations is proportional to the squared fitness effect of deleterious alleles, multiplied by the squared mean number of mutations per chromosome, yielding (*sh*)^2^ *×* (*U/sh*)^2^ = *U* ^2^ in the case of deleterious mutations (where *U* is the deleterious mutation rate per chromosome and *sh* the heterozygous fitness effect of mutations), and *u*^2^ *×* (*f n/*2)^2^ in the present model. Although the effective population size may be reduced by background selection effects, *N*_e_ should stay relatively constant along the chromosome as long as *R* is not too small (given that the model assumes a uniform distribution of crossovers along the chromosome), in which case the effect of segregating TEs on *N*_e_ is approximately *N*_e_ *≈ N* exp(*−uf n/R*) (Hudson and Kaplan, 1995; Charlesworth, 1996).

Figure 7 shows the evolutionarily stable chromosome map length *R*_ES_ for different numbers of TE families *f* and for parameter values leading to *n ≈* 10 (per family) at the deterministic equilibrium. Figure 7A shows *R*_ES_ as a function of the transposition rate *u*, the parameter *θ* measuring the direct fitness cost of ectopic recombination being set to zero, but assuming an inherent fitness cost *c* = 0.001 per crossover (in the absence of cost, the average map length fluctuates widely in the simulations when *u* is small, due to the fact that indirect selection becomes very weak as *R* increases). Figure 7B shows *R*_ES_ as a function of the cost of ectopic recombination *θ*, the inherent cost *c* being set to zero. In the case of a single TE family (*f* = 1), the Hill-Robertson effect is not sufficiently strong (relative to direct selection and to the deterministic component of indirect selection) to maintain recombination: *R* is predicted to evolve towards zero as this minimizes ectopic recombination (when *θ >* 0) or any inherent cost of recombination (when *c >* 0) and allows a better purging of elements in positive LD. As shown by Figure 7, this is confirmed by the simulation results. The Hill-Robertson effect (generating negative LD and favoring recombination) becomes more important when larger numbers of elements are segregating. Equation 21 predicts that for a given load of transposons (fixed *n*) the strength of selection for recombination due to the Hill-Robertson effect increases with the transposition rate *u*, which is confirmed by Figure 7A. For low values of *u* (and low *R*_ES_), important fluctuations of the mean chromosome map length *R* and of the mean number of elements per genome and per family *n* can be observed in the simulations (see Figure S5 for an example): *n* increases rapidly when *R* reaches low values due to the decreased efficiency of selection caused by the Hill-Robertson effect, but the increase in *n* then favors higher recombination rates, leading to more efficient selection against TEs and to a decrease in *n*. For *u* = 0.01, the approximations shown above predict values of *R*_ES_ in the range 0.1 – 0.4 with 10 or 20 TE families even when the cost of ectopic recombination is high (green and red curves in Figure 7B). However, while the simulations agree with the analytical results when the cost of ectopic recombination is weak, above a threshold value of *θ* (to the right of the right-most green and red dots on Figure 7B) *R*_ES_ becomes too small to maintain TEs at a stable equilibrium, and the number of TEs per individual quickly reaches very high values. This threshold is reached sooner (*i.e.*, at lower values of *θ*) when *Nu* is smaller (for a fixed *n* at deterministic equilibrium) or when *n* is higher (results not shown). Below the threshold, the discrepancy observed between the model predictions and the simulations as *θ* increases is possibly due to the interaction between the cost of recombination and the strength of selection against TEs, which is not taken into account in Roze’s (2021) analysis of selection for recombination due to the Hill-Robertson effect. Overall, Figure 7B shows that assuming substantial direct costs of recombination (due to a positive correlation between *R* and the rate of ectopic recombination) generally drives the evolution of *R* below the minimal value at which TEs can be maintained at a stable equilibrium.

**Figure 7.**
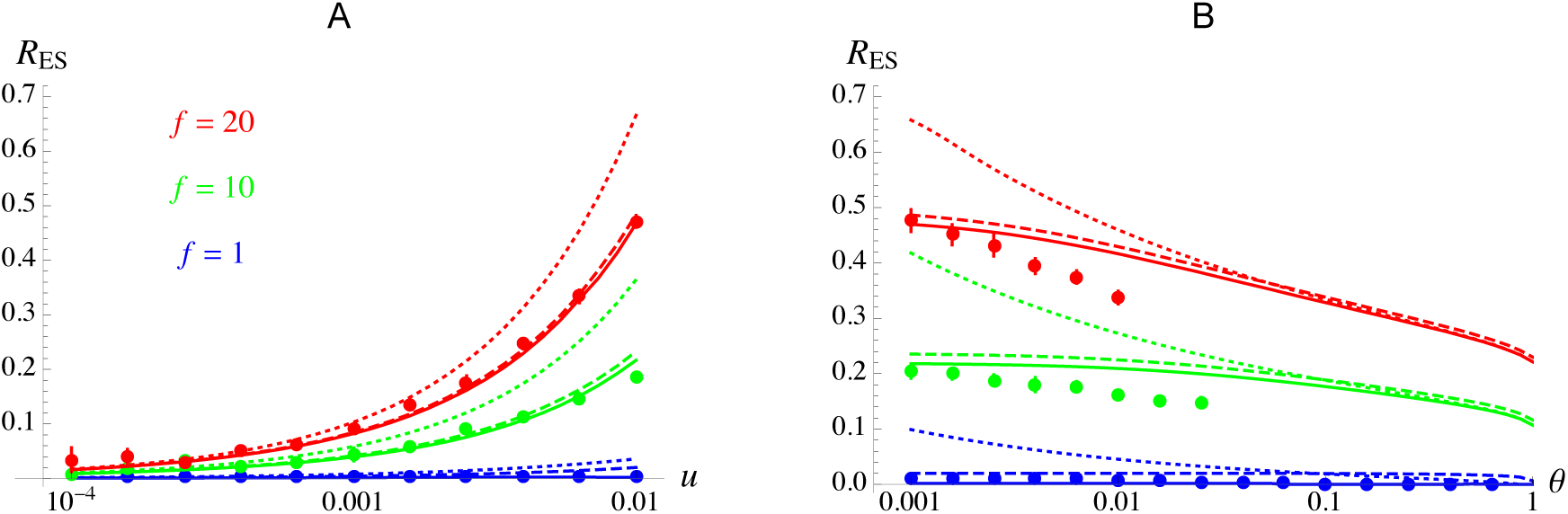
Evolutionarily stable chromosome map length *R*_ES_ (in Morgans) for different numbers of segregating TE families *f* . A: as a function of the transposition rate *u* (on a log scale), in the absence of cost of ectopic recombination (*θ* = 0), but with an inherent cost of crossovers set to *c* = 0.001. B: as a function of the cost of ectopic recombination *θ* (on a log scale), for *c* = 0. Solid curves correspond to analytical predictions, obtained by solving *s*_dir_ + *s*_ind,det_ + *s*_ind,HR_ = 0 for *R*, where *s*_dir_ is given by equation 19, *s*_ind,det_ by equation 20 and *s*_ind,HR_ by equation A114 in the Supplementary Methods. Dashed curves correspond to analytical predictions obtained when *s*_ind,HR_ is approximated by equation 21, and dotted curves to the predictions obtained when ignoring the deterministic component of indirect selection (obtained by solving *s*_dir_ + *s*_ind,HR_ = 0 for *R*). Dots: simulation results. On the right of the right-most points in B, TEs are accumulating (no stable equilibrium) and the simulation has to be stopped. Parameter values: *N* = 10^5^, *u* = 0.01 (in B), *v* = *u/*100, *α* = 0, *n* = 10 per family — in the simulations shown in B, the value of *β*^°^(defined in equation 10) is set so that *n* = 10 (using equation 11 for *n*) when *R* is at its predicted evolutionarily stable value; in A, *β*^°^ = *β* = *u/*10.

## DISCUSSION

Theoretical models developed in the early 1980s identified two main mechanisms that may limit the spread of TEs within their host genomes (Charlesworth and Charlesworth, 1983; Langley et al., 1983; Kelleher et al., 2020): copy-number-dependent transposition (lower rates of transposition as the number of elements increase), and synergistic purifying selection. While transcriptional and post-transcriptional silencing of TEs correspond to plausible mechanisms generating copy-number-dependent transposition (Deniz et al., 2019; Kelleher et al., 2020; Almeida et al., 2022), ectopic recombination has been identified as a possible source of synergistic epistasis among elements (Montgomery et al., 1987; Langley et al., 1988). Indirect evidence suggests that ectopic recombination likely plays an important role in the containment of TEs (Bartolomé et al., 2002; Song and Boissinot, 2007; Petrov et al., 2011; Bonchev and Willi, 2018); however, to what extent this process generates synergistic selection against TE copies remains difficult to test. Recently, Lee (2022) used population genomic data from *D. melanogaster* to infer the degree of synergism among TEs from patterns of linkage disequilibrium between elements, based on the assumption that synergistic (*i.e.*, negative) epistasis generates negative LD. The results showed positive LD between tightly linked TEs, but negative LD between more distant TEs. While positive LD may have been generated by admixture among divergent populations (Sohail et al., 2017; Sandler et al., 2021) or by the fact that the analysis was restricted to TE insertions present in a small number of individuals (Good, 2022), negative LD among distant TEs was interpreted as evidence for synergistic epistasis.

The present paper shows that classical results on the effect of epistasis on LD between deleterious mutations cannot be directly transposed to TEs, however, due to the fact that the transposition process is a source of positive LD among elements. Under the assumptions of the classical model of synergistic purifying selection against TEs used in this paper (Charlesworth, 1991; Dolgin and Charlesworth, 2006, 2008), the effect of transposition is stronger than the effect of negative epistasis and generates positive LD at equilibrium. Note that this result holds for both class 1 and class 2 elements (Wells and Feschotte, 2020): in the case of class 2 elements (DNA transposons), the simple movement of TEs through cut-and-paste mechanisms does not generate any LD when the original insertion is excised, but positive LD is generated (on average) when the original insertion is retained and the copy number increases. Strikingly, in an infinite population the LD between loosely linked loci (*r_ij_ » u*) scaled by the product of genetic variances at both loci (*D_ij_/p_i_q_i_p_j_q_j_*) is approximately *β/r_ij_*where *β* is the strength of negative epistasis (from equation 16, assuming *v*, *α « u*), and thus exactly opposite to the result obtained in the case of deleterious mutations (*−β/r_ij_*, e.g., Barton, 1995). Negative LD may be generated by the Hill-Robertson effect in finite populations, however, the relative importance of this effect increasing with the degree of linkage among loci. In a large population with frequent recombination, the sum of all LD among TEs is predicted to be slightly positive, the distribution of the number of TEs per genome staying very close to a Poisson distribution. Within the genome, LD between closely linked insertion sites may be either positive or negative depending on the relative importance of deterministic and stochastic effects (Figure 5).

These predictions seem at odds with the observation of negative LD between insertions present on different chromosomes of *D. melanogaster* (Lee, 2022). Negative LD between loosely linked loci could arise due to the Hill-Robertson effect in the present model, but only in the case of a very low effective population size, which seems unlikely given that the source population shows high genetic diversity and very low spatial structure (Lack et al., 2015). Epistasis may also generate negative LD despite the positive effect of transposition under some forms of fitness functions, but Figure 2 indicates that the curvature of the fitness function has to be quite strong for the effect of epistasis to take over. Alternatively, the discrepancy between theoretical and empirical results raises the possibility that the model considered here does not provide an adequate representation of transposon dynamics in natural populations. Various extensions of the model would be worth exploring, introducing for example different probabilities of ectopic recombination between pairs of elements separated by different genomic distances, or insertion biases of new TE copies. Furthermore, the present model does not include any mechanism of host regulation of transposition. TE silencing mechanisms have been increasingly well described over recent years; in particular, the system based on Piwi-interacting RNAs (which inactivates TEs once an element has inserted into a piRNA cluster) is present in a variety of organisms and generates a form of copy-number-dependent transposition (Brennecke et al., 2007; Gunawardane et al., 2007; Kelleher et al., 2020). Several models of TE dynamics incorporating silencing through piRNAs have been proposed recently (Kelleher et al., 2018; Kofler, 2019, 2020; Kofler et al., 2022; Tomar et al., 2022), and it would be interesting to explore how host regulation may affect LD patterns among TEs using this type of models.

From a more general perspective, the present results help to reconcile two different views on the effect of reproductive systems and recombination on TE dynamics: on one hand, sexual reproduction is assumed to favor the spread of TEs (Hickey, 1982) while on the other hand, the lack of recombination may lead to TE accumulation through Muller’s ratchet (Arkhipova and Meselson, 2005a; Dolgin and Charlesworth, 2008). When population size is large and recombination is frequent, the Hill-Robertson effect among TEs often stays negligible and the main effect of recombination is to decrease the excess variance in TE number caused by the transposition process, thus helping the spread of TEs (in agreement with Hickey’s view). However, the Hill-Robertson effect may predominate when effective population size is small and/or recombination is rare, in particular when the number of TEs per genome is large: in this case, recombination increases the effect of selection against TEs. This also sheds light on the simulation results obtained by Dolgin and Charlesworth (2006), showing that a transition towards asexuality may either lead to an accumulation of TEs or to their elimination, the second outcome occurring above a threshold population size (and requiring TE excision if all individuals are initially loaded with TEs). To date, empirical comparisons between sexual and asexual species have not shown any clear trend. While the first TE surveys in the putatively ancient asexual bdelloid rotifers suggested a low TE content (Arkhipova and Meselson, 2000, 2005b), more recent work has shown a similar level of TE content and activity to what is observed in sexual rotifer species (Nowell et al., 2021). Similarly, comparisons of closely related sexual and asexual arthropod species showed little difference in TE content (Bast et al., 2016; Jaron et al., 2022). However, it is possible that extant asexual species originated from sexual ancestors in which the level of TE activity was low, while new asexual lineages in which TE activity is high tend to go extinct due to TE accumulation (Arkhipova and Meselson, 2005a), and it would thus be of interest to explore TE dynamics in newly derived asexual lineages. Transitions towards self-fertilization may also either lead to an increased or decreased efficiency of selection against TEs depending on the relative importance of deterministic and stochastic forces, with the additional effect that homozygosity may decrease the rate of ectopic recombination among elements (Montgomery et al., 1991; Wright and Schoen, 1999; Morgan, 2001; Bonchev and Willi, 2018). Measuring LD among TEs in selfing or asexual populations (and contrasting LD within and between TE families) may shed further light on the selective forces acting on TEs, and should be made easier by the recent progress in long-reads sequencing technologies.

Linkage disequilibrium between selected sites is the source of indirect selection on recombination modifier alleles (Felsenstein, 1974). In classical models of deleterious mutations, synergistic epistasis generates negative LD, causing recombination to decrease the mean fitness of offspring, but increase the variance in fitness and the efficiency of selection. The benefit of increasing the variance in fitness becomes stronger than the short-term cost of reducing mean fitness as recombination decreases, so that non-zero rates of recombination are predicted to be maintained at equilibrium (Charlesworth, 1990; Barton, 1995; Charlesworth and Barton, 1996). In the deterministic regime of the present model, positive LD among TEs is maintained despite synergistic epistasis and recombination therefore has opposite effects, increasing the mean fitness of offspring but decreasing the variance in fitness. The disadvantage caused by the reduced efficiency of selection often predominates (in particular when the number of TEs per genome is large), disfavoring recombination. However, interference effects caused by finite population size favor recombination and tend to become stronger than deterministic effects as recombination decreases: in the absence of any direct fitness cost of recombination, this may favor the maintenance of high crossover rates when the number of TEs per chromosome is large. It seems likely that ectopic recombination among TEs will generate a direct fitness cost for recombination, however, and in the present model, this direct cost tends to drive the crossover rate towards values at which TEs cannot be maintained at a stable equilibrium anymore. Here again, using models of TE bursts followed by silencing through host regulation would be of interest, as a burst of TEs may transiently favor or disfavor recombination depending on the balance between the direct cost of ectopic recombination and interference effects. The joint evolution of local TE density and local recombination rate, or the effect of interactions between TEs and classical deleterious mutations may also be worth exploring. From an empirical perspective, more detailed knowledge on ectopic recombination among TEs is needed in order to better assess its potential consequences: in particular, to what extent it correlates with meiotic recombination rates, whether it may occur among silenced TEs, and how the physical distance among TEs may affect the probability of ectopic exchange. More generally, assessing the relative contribution of segregating TEs to the variance in fitness within populations would help us to better understand the evolutionary consequences of our load of selfish DNA.

**Data availability.** Derivations are provided in the *Mathematica* notebook available as Supplementary Material at doi.org/10.5281/zenodo.7233938, along with the C++ codes used to run the simulations.

## Acknowledgements

**Acknowledgements.** I thank the Bioinformatics and Computing Service of Roscoff’s Biological Station (Abims platform) for computing time, and Arnaud Le Rouzic, Vincent Castric and three anonymous reviewers for useful comments.

**Funding.** This work was funded by the Agence Nationale pour la Recherche (SelfRecomb project: ANR-18-CE02-0017-02, and GenAsex project: ANR-17-CE02-0016-01).

**Competing interests.** None declared.

