## Supplementary Figures for "Causes and consequences of linkage disequilibrium among transposable elements within eukaryotic genomes"

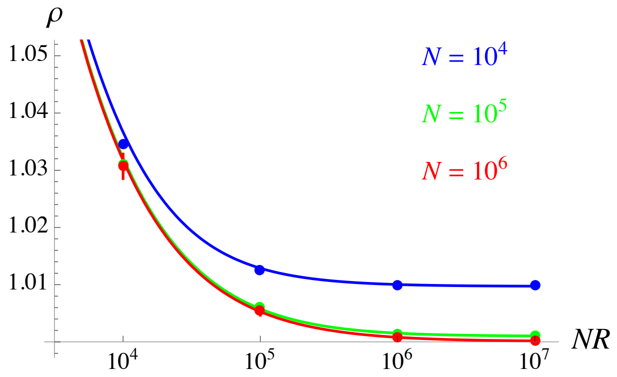


**Figure S1.** Zoom on the right part of Figure 1 for $Nu = 100$ and $\overline{n} = 10$, where the curves correspond to the more accurate expression of $\mathcal{E}_1$ for high $R$ obtained by replacing $x$ by $\left(1 - e^{-2x}\right)/2$ in the denominator of equation A38 of the Supplementary Methods. Approximation A39 for $\mathcal{E}_1$ (which assumes that $R$ is not too large, and corresponds to the black curve in Figure 1) is indistinguishable from the red curve.

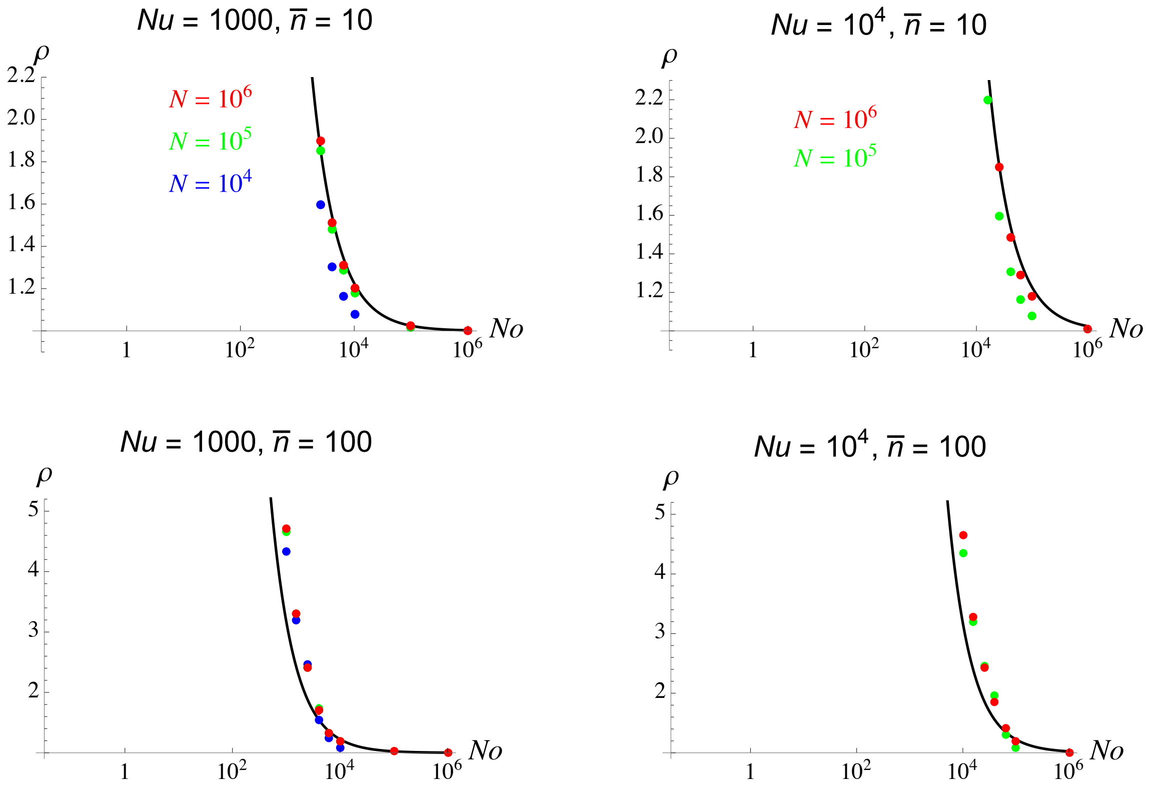


**Figure S2.** Same as Figure 4 in the main text, with $Nu = 1000$ (left) and $Nu = 10^4$ (right). To the left of the left-most points, TEs are eliminated from the population during the simulations when $\overline{n} = 10$ (top figures) and when $\overline{n} = 100$ and $n_{\text{init}} = 10$ (bottom figures), while they accumulate when $\overline{n} = 100$ and $n_{\text{init}} = 100$.

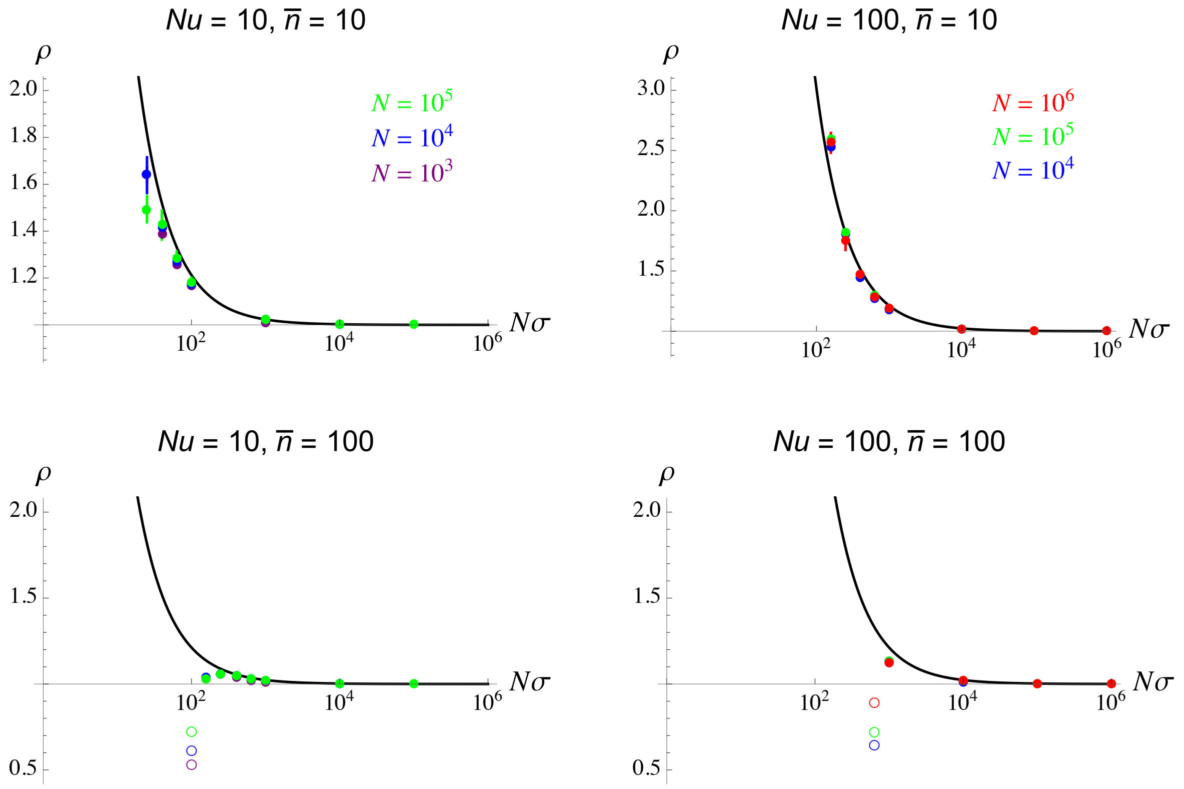


**Figure S3.** Effect of genetic associations among TEs at different sites on the variance in TE number per individual ($\rho$), as a function of the product of population size $N$ and the rate of sex $\sigma$ (on a log scale) in a facultatively sexual population, for different values of $Nu$ and $\overline{n}$ (the mean number of TEs per individual at the deterministic equilibrium, given by equation 10 in the main text). Curves correspond to the deterministic approximation given by equation A49 in the Supplementary Methods). Dots: simulation results; the different colors correspond to different values of population size $N$. On the left of the left-most filled circles of each figure, TEs are either eliminated from the population during the simulation or keep accumulating: TEs are eliminated in the top figures ($\overline{n} = 10$), but accumulate in the bottom figures ($\overline{n} = 100$; as in Figures 1 and 4 in the main text, empty circles correspond to averages over the last 10 data points of the simulations). Parameter values: $R = 10$, $v = u/100$, $\alpha = 0$, $\beta = u/10$ (top figures) or $u/100$ (bottom figures).

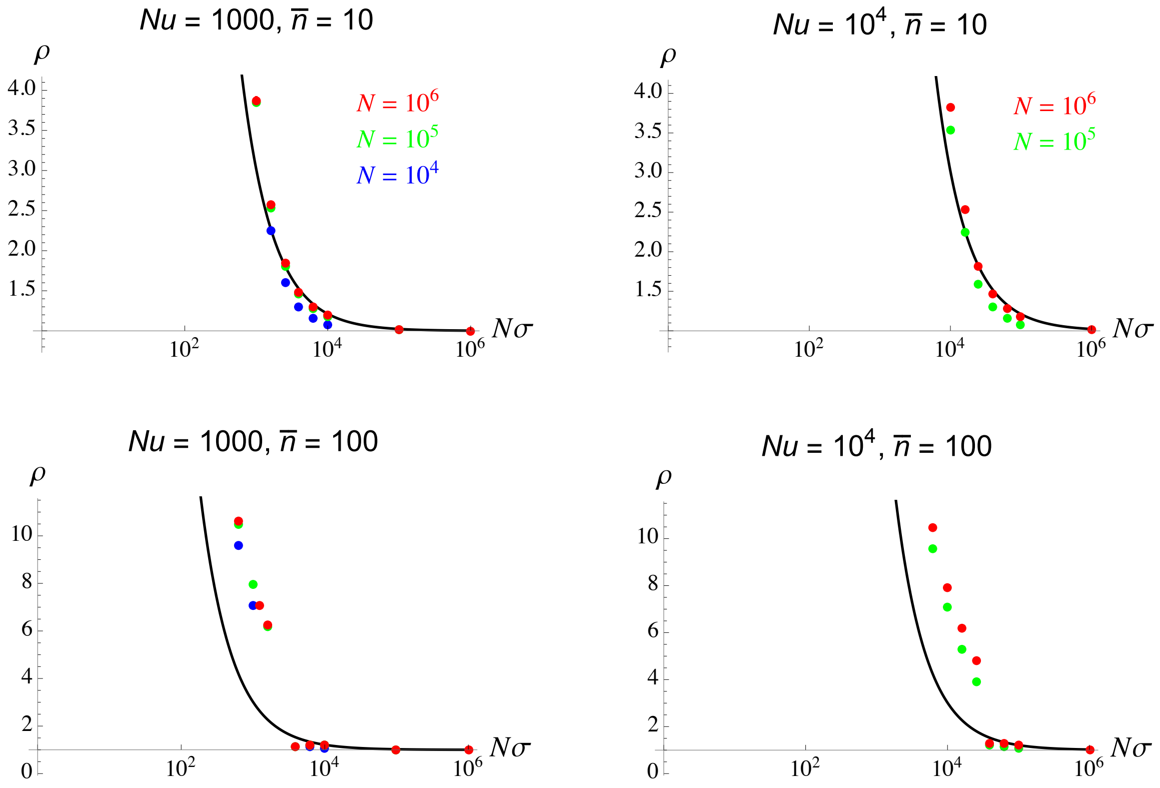


**Figure S4.** Same as Figure S3 with $Nu = 1000$ (left) and $Nu = 10^4$ (right). To the left of the left-most points, TEs are eliminated from the population during the simulations when $\overline{n} = 10$ (top figures) and when $\overline{n} = 100$ and $n_{\text{init}} = 10$ (bottom figures), while they accumulate when $\overline{n} = 100$ and $n_{\text{init}} = 100$.

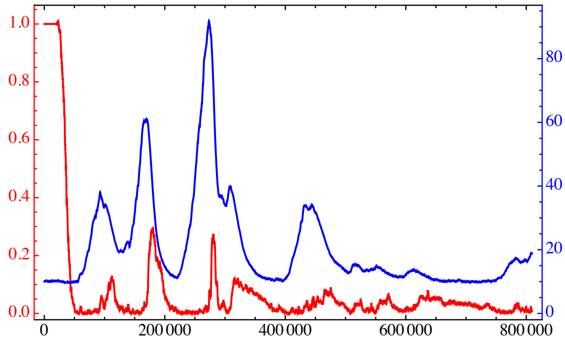


**Figure S5.** Average chromosome map length $\overline{R}$ (in red) and average number of elements per genome and per TE family (in blue) over the course of a simulation including a recombination modifier affecting $R$ (the x-axis shows time in generations, $R$ is fixed to 1 during the first 20,000 generations). Parameter values are those of Figure 7A with $f = 20$ and $u = 1.58 \times 10^{-4}$.
