## Supplementary Methods for "Causes and consequences of linkage disequilibrium among transposable elements within eukaryotic genomes"

**1. Notation.** Genetic associations among TEs are defined using the multilocus notation of Barton and Turelli (1991) and Kirkpatrick et al. (2002). Throughout the following, $X_{i,\varnothing}$ and $X_{\varnothing,i}$ correspond to indicator variables that equal 1 if a TE copy is present at insertion site $i$ on the maternally and paternally inherited haplotypes of an individual (respectively), and 0 otherwise. As the model does not include any sex-of-origin effect, we have $\mathrm{E}[X_{i,\varnothing}] = \mathrm{E}[X_{\varnothing,i}]$ (where E stands for the average over all individuals in the population) corresponding to the frequency of insertions at site $i$ in the population, denoted $p_i$ and supposed to be small. Using the centered variables $\zeta_{i,\varnothing} = X_{i,\varnothing} - p_i$ and $\zeta_{\varnothing,i} = X_{\varnothing,i} - p_i$, genetic associations are defined as:

$$D_{\mathbb{U},\mathbb{V}} = \mathrm{E}[\zeta_{\mathbb{U},\mathbb{V}}] \tag{A1}$$

with:

$$\zeta_{\mathbb{U},\mathbb{V}} = \left(\prod_{i\in\mathbb{U}} \zeta_{i,\varnothing}\right)\left(\prod_{j\in\mathbb{V}} \zeta_{\varnothing,j}\right) \tag{A2}$$

and where $\mathbb{U}$ and $\mathbb{V}$ represent sets of insertions sites. Because the model assumes no difference between sexes, we have $D_{\mathbb{U},\mathbb{V}} = D_{\mathbb{V},\mathbb{U}}$. For simplicity, associations between TEs present on the same haplotype ($D_{\mathbb{U},\varnothing} = D_{\varnothing,\mathbb{U}}$) will be denoted $D_{\mathbb{U}}$: in particular, $D_{ij}$ corresponds to the linkage disequilibrium between insertions at sites $i$ and $j$. Using the fact that $X_{i,\varnothing}^2 = X_{i,\varnothing}$ and $X_{\varnothing,i}^2 = X_{\varnothing,i}$ (since these variables equal 0 or 1), eliminated indices that may appear in associations can be eliminated using the relation given by equation 5 in Kirkpatrick et al. (2002):

$$D_{\mathbb{U}ii,\mathbb{V}} = p_i q_i D_{\mathbb{U},\mathbb{V}} + (1 - 2p_i)\, D_{\mathbb{U}i,\mathbb{V}} \tag{A3}$$

where $\mathbb{U}$ and $\mathbb{V}$ may be any set of loci and $q_i = 1 - p_i$; for example, $D_{ii,\varnothing} = p_i q_i$, while $D_{iij,\varnothing} = (1 - 2p_i)\, D_{ij,\emptyset}$ (since $D_{i,\varnothing} = D_{\varnothing,i} = 0$, while $D_{\varnothing} = 1$). The derivations presented here assume that the parameters $u$, $v$, $\alpha$ and $\beta$ are small (of order $\epsilon$, where $\epsilon$ is a scaling factor); furthermore, recombination rates are assumed to be high, so that genetic associations between insertions at different sites remain small (of order $\epsilon$); however, we will see that some of the results can be extended to tight linkage among insertions. Variables measured after transposition, excision and selection will be denoted with superscripts 't', 'e' and 's', respectively, while variables measured at the next generation (after reproduction) will be denoted with a prime.

**2. Mean number of TEs per genome.** The average number of TEs per genome, $\overline{n} = 2\sum_i p_i$ changes during transposition/excision and during selection. Using the notation defined above, and noting that the number of TEs in an individual is given by $n = \sum_i (X_{i,\varnothing} + X_{\varnothing,i})$, the fitness of an individual can be written as:

$$\begin{aligned} W &= \exp\left[-\alpha \sum_i (X_{i,\varnothing} + X_{\varnothing,i}) - \frac{\beta}{2} \sum_{i \neq j} (X_{i,\varnothing} + X_{\varnothing,i})(X_{j,\varnothing} + X_{\varnothing,j})\right] \\ &= 1 - \alpha\,\overline{n} - \frac{\beta}{2}\overline{n}^2 - (\alpha + \beta\overline{n}) \sum_i (\zeta_{i,\varnothing} + \zeta_{\varnothing,i}) \\ &\quad - \frac{\beta}{2} \sum_{i \neq j} (\zeta_{ij,\varnothing} + \zeta_{\varnothing,ij} + \zeta_{i,j} + \zeta_{j,i}) + o(\epsilon) \end{aligned} \tag{A4}$$

where the last sum is over all pairs of insertion sites $i$ and $j$ (each pair being counted twice in the sum). From this, the average fitness in the population is:

$$\overline{W} = 1 - \alpha\,\overline{n} - \frac{\beta}{2}\overline{n}^2 - \beta \sum_{i \neq j} (D_{ij} + D_{i,j}) + o(\epsilon)\,, \tag{A5}$$

leading to:

$$\begin{aligned} \frac{W}{\overline{W}} &= 1 - \sum_i a_i (\zeta_{i,\varnothing} + \zeta_{\varnothing,i}) \\ &\quad - \sum_{i \neq j} a_{ij} (\zeta_{ij,\varnothing} + \zeta_{\varnothing,ij} - 2D_{ij} + \zeta_{i,j} + \zeta_{j,i} - 2D_{i,j}) + o(\epsilon^2) \end{aligned} \tag{A6}$$

with $a_i = \alpha + \beta\overline{n}$ and $a_{ij} = \beta/2$. The effect of selection on the frequency of insertions at site $i$ is given by:

$$\Delta_{\text{sel}} p_i = \mathrm{E}\left[\frac{W}{\overline{W}} \frac{X_{i,\varnothing} + X_{\varnothing,i}}{2}\right] - p_i = \mathrm{E}\left[\frac{W}{\overline{W}} \frac{\zeta_{i,\varnothing} + \zeta_{\varnothing,i}}{2}\right]. \tag{A7}$$

From equation A6, and using the fact that $D_{i,j} = 0$, $D_{ij,k} = 0$ under random mating, this is:

$$\Delta_{\text{sel}} p_i = -(\alpha + \beta\overline{n}) \sum_j D_{ij} - \frac{\beta}{2} \sum_{j \neq k} D_{ijk} + o(\epsilon)\,. \tag{A8}$$

Neglecting linkage disequilibria and using $D_{ii} = p_i q_i$ (from equation A3), we have to the first order in $p_i$ and in $\epsilon$:

$$\Delta_{\text{sel}} p_i \approx -(\alpha + \beta\overline{n})\, p_i, \tag{A9}$$

leading to:

$$\Delta_{\mathrm{sel}}\overline{n} \approx -\left(\alpha + \beta\overline{n}\right)\overline{n}\,. \tag{A10}$$

Furthermore, the changes in $\overline{n}$ due to transposition and excision are given by:

$$\Delta_{\mathrm{transp}}\overline{n} = u\overline{n}, \qquad \Delta_{\mathrm{exc}}\overline{n} = -v\overline{n} \tag{A11}$$

so that the equilibrium value of $\overline{n}$ (obtained by solving $\Delta_{\mathrm{sel}}\overline{n} + \Delta_{\mathrm{transp}}\overline{n} + \Delta_{\mathrm{exc}}\overline{n} = 0$) is given by:

$$\overline{n} \approx \frac{u - v - \alpha}{\beta} \tag{A12}$$

(Charlesworth, 1991). Note that from equation A12, the effective strength of selection acting against each TE copy (the coefficient $a_i = \alpha + \beta\overline{n}$ in equation A6) is approximately $u - v$.

**3. Linkage disequilibrium between TEs.** The linkage disequilibrium between TE insertions at sites $i$ and $j$ at the next generation, is given by:

$$D_{ij}{}' = \left(1 - r_{ij}\right) D_{ij}{}^{\mathrm{s}} + r_{ij}\, D_{i,j}{}^{\mathrm{s}} \tag{A13}$$

where $r_{ij}$ is the recombination rate between sites $i$ and $j$ and where $D_{ij}{}^{\mathrm{s}}$, $D_{i,j}{}^{\mathrm{s}}$ are measured after selection (before reproduction). The change in $D_{ij}$ caused by selection is given by (e.g., Barton and Turelli, 1991):

$$D_{ij}{}^{\mathrm{s}} = \mathrm{E}\left[\frac{W}{\overline{W}}\frac{\zeta_{ij,\varnothing} + \zeta_{\varnothing,ij}}{2}\right] - \Delta_{\mathrm{sel}}p_i\, \Delta_{\mathrm{sel}}p_j. \tag{A14}$$

From equation A6 and A9, and using the fact that $D_{i,j} = 0$, $D_{ij,k} = 0$, $D_{ijk,l} = 0$ under random mating, this is:

$$D_{ij}{}^{\mathrm{s}} = D_{ij}{}^{\mathrm{t}} - \left(\alpha + \beta\overline{n}\right)\sum_{k} D_{ijk}{}^{\mathrm{t}} - \frac{\beta}{2}\sum_{k \neq l}\left(D_{ijkl}{}^{\mathrm{t}} + D_{ij,kl}{}^{\mathrm{t}} - 2D_{ij}{}^{\mathrm{t}}D_{kl}{}^{\mathrm{t}}\right) + o(\epsilon)\,. \tag{A15}$$

where $D_{ijk}{}^{\mathrm{t}}$, $D_{ijkl}{}^{\mathrm{t}}$ and $D_{ij,kl}{}^{\mathrm{t}}$ are three and four-locus associations defined by equation A1. Assuming that linkage disequilibria are of order $\epsilon$, only the terms for $k = i$, $l = j$ and $k = j$, $l = i$ in the second sum remain. Using the fact that $D_{iijj} = p_i q_i p_j q_j + \left(1 - 2p_i\right)\left(1 - 2p_i\right) D_{ij}$ (from equation A3), one obtains:

$$D_{ij}{}^{\mathrm{s}} \approx D_{ij}{}^{\mathrm{t}} - \beta\, p_i p_j\,. \tag{A16}$$

Similarly, we have:

$$D_{i,j}{}^{\mathrm{s}} \approx D_{i,j}{}^{\mathrm{t}} - \beta\, p_i p_j \,. \tag{A17}$$

In order to compute the effect of transposition on $D_{ij}$, we first consider that the number of possible insertions sites $L$ is finite (but much larger than the number of TEs present in the genome), and then take the limit as $L$ tends to infinity. $D_{ij,\varnothing}$ after transposition is given by:

$$D_{ij,\varnothing}{}^{\mathrm{t}} = \mathrm{E}\left[\left(X_{i,\varnothing}{}^{\mathrm{t}} - p_i{}^{\mathrm{t}}\right)\left(X_{j,\varnothing}{}^{\mathrm{t}} - p_j{}^{\mathrm{t}}\right)\right] \tag{A18}$$

where

$$X_{i,\varnothing}{}^{\mathrm{t}} = X_{i,\varnothing}{}^{\mathrm{e}} + \left(1 - X_{i,\varnothing}{}^{\mathrm{e}}\right)\frac{u}{2L}\sum_{k\neq i}\left(X_{k,\varnothing}{}^{\mathrm{e}} + X_{\varnothing,k}{}^{\mathrm{e}}\right). \tag{A19}$$

Indeed, a TE is present on the maternally inherited chromosome at site $i$ either if it was already present before transposition ($X_{i,\varnothing}{}^{\mathrm{e}} = 1$, first term of equation A19), or if one of the new $u\,n = u\sum_{k\neq i}\left(X_{k,\varnothing}{}^{\mathrm{e}} + X_{\varnothing,k}{}^{\mathrm{e}}\right)$ copies inserts at this site, which happens with probability $1/(2L)$ (second term of equation A19). Dropping e superscripts for clarity, replacing $X_{i,\varnothing}$ by $\zeta_{i,\varnothing} + p_i$, and assuming that LD and $p_i$ are small yields $p_i{}^{\mathrm{t}} = \mathrm{E}\left[X_{i,\varnothing}{}^{\mathrm{t}}\right] \approx p_i + \frac{u}{2L}\,\overline{n}$, while:

$$\begin{aligned} D_{ij,\varnothing}{}^{\mathrm{t}} \approx \mathrm{E}\Bigg[&\left[\zeta_{i,\varnothing}\left(1 - \frac{u}{2L}\,\overline{n}\right) + \frac{u}{2L}\sum_{k\neq i}\left(\zeta_{k,\varnothing} + \zeta_{\varnothing,k} - \zeta_{ik,\varnothing} - \zeta_{i,k}\right)\right] \\ &\times\left[\zeta_{j,\varnothing}\left(1 - \frac{u}{2L}\,\overline{n}\right) + \frac{u}{2L}\sum_{k\neq j}\left(\zeta_{k,\varnothing} + \zeta_{\varnothing,k} - \zeta_{jk,\varnothing} - \zeta_{j,k}\right)\right]\Bigg]. \end{aligned} \tag{A20}$$

Assuming that linkage disequilibria are of order $\epsilon$, this yields:

$$D_{ij}{}^{\mathrm{t}} \approx D_{ij}{}^{\mathrm{e}} + \frac{u}{2L}\left(p_i + p_j\right). \tag{A21}$$

The term $D_{ij}{}^{\mathrm{e}}$ in equation A21 stems from the product of $\zeta_{i,\varnothing}$ on the first line of equation A20 with $\zeta_{j,\varnothing}$ on the second line, while the term in $p_i$ stems from the product of $\zeta_{i,\varnothing}$ on the first line with the term for $k = i$ of the sum on the second line, yielding $u/(2L)\left(D_{ii,\varnothing} + D_{i,i} - D_{iij,\varnothing} - D_{ij,i}\right)$, where $D_{ii,\varnothing} = p_i q_i \approx p_i$, while $D_{i,i} = 0$, $D_{ij,i} = 0$ under random mating and $D_{iij,\varnothing} = (1 - 2p_i)\, D_{ij}$ is of order $\epsilon$. Similarly, the product of $\zeta_{j,\varnothing}$ on the second line of equation A20 with the term for $k = j$ of the sum on the first line yields the term in $p_j$ in equation A21. All other terms of equation A20 are $o(\epsilon)$. The same reasoning yields:

$$D_{i,j}{}^{\mathrm{t}} \approx D_{i,j}{}^{\mathrm{e}} + \frac{u}{2L}\left(p_i + p_j\right). \tag{A22}$$

The effect of TE excision is obtained similarly. We have:

$$D_{ij,\varnothing}{}^{\mathrm{e}} = \mathrm{E}\left[\left(X_{i,\varnothing}{}^{\mathrm{e}} - p_i{}^{\mathrm{e}}\right)\left(X_{j,\varnothing}{}^{\mathrm{e}} - p_j{}^{\mathrm{e}}\right)\right] \tag{A23}$$

with $X_{i,\varnothing}{}^{\mathrm{e}} = (1-v)\,X_{i,\varnothing}$, $p_i{}^{\mathrm{e}} = (1-v)\,p_i$, yielding:

$$D_{ij}{}^{\mathrm{e}} = (1-v)^2\,D_{ij} \approx D_{ij}, \quad D_{i,j}{}^{\mathrm{e}} \approx (1-v)^2\,D_{i,j} = 0. \tag{A24}$$

Altogether, equations A13, A16, A17, A21, A22 and A24 yield, at equilibrium:

$$D_{ij} \approx \frac{1}{r_{ij}}\left[\frac{u}{2L}\,(p_i + p_j) - \beta\,p_i p_j\right]. \tag{A25}$$

The effect of LD on the variance in the number of TEs per genome is twice the sum of linkage disequilibria over all pairs of possible insertion sites, and is thus given by:

$$2\sum_{i\neq j} D_{ij} \approx \frac{1}{r_{\mathrm{h}}}\left[u\overline{n} - \beta\frac{\overline{n}^2}{2}\right], \tag{A26}$$

where $r_{\mathrm{h}}$ is the harmonic mean recombination rate between all possible pairs of insertion sites. Using equation A12, this yields:

$$2\sum_{i\neq j} D_{ij} \approx \frac{\overline{n}}{2r_{\mathrm{h}}}\,(u + v + \alpha)\,. \tag{A27}$$

Equation A27 shows that linkage disequilibria are positive due to the effect of transposition, and thus tend to increase the variance in the number of TEs per genome. However, equation A25 diverges as the recombination rate $r_{ij}$ tends to zero, and $r_{\mathrm{h}}$ thus cannot be computed in the case of a continuous genetic map. Following Roze (2014, 2016, 2021), more accurate results can be obtained in the case of tightly linked loci by relaxing the assumption that linkage disequilibria are small during the derivation of recurrence equations. Using the fact that $D_{iij} = (1-2p_i)\,D_{ij} \approx D_{ij}$ and still neglecting associations between more than two loci, equation A15 then yields:

$$D_{ij}{}^{\mathrm{s}} \approx \left[1 - 2\left(\alpha + \beta\overline{n}\right)\right] D_{ij}{}^{\mathrm{t}} - \beta\,p_i p_j\,. \tag{A28}$$

Note that the term in $-\beta\,D_{iijj}$ in equation A15 also generates a term in $-\beta\,D_{ij}{}^{\mathrm{t}}$, but throughout the following we will generally consider that $\overline{n}$ is large, so that this term should be negligible relative to the term $-2\beta\overline{n}D_{ij}{}^{\mathrm{t}}$ that appears in equation A28. Similarly, we have:

$$D_{i,j}{}^{\mathrm{s}} \approx \left[1 - 2\left(\alpha + \beta\overline{n}\right)\right] D_{i,j}{}^{\mathrm{t}} - \beta\,p_i p_j\,. \tag{A29}$$

Furthermore, equation A20 now yields (still neglecting associations between more than two loci):

$$D_{ij}{}^{\mathrm{t}} \approx \left(1 - \frac{u}{2L}\,\overline{n}\right) D_{ij}{}^{\mathrm{e}} + \frac{u}{2L}\left(p_i + p_j + \sum_k D_{ik} + \sum_k D_{jk}\right) \tag{A30}$$

and similarly

$$D_{i,j}{}^{\mathrm{t}} \approx \left(1 - \frac{u}{2L}\,\overline{n}\right) D_{i,j}{}^{\mathrm{e}} + \frac{u}{2L}\left(p_i + p_j + \sum_k D_{ik} + \sum_k D_{jk}\right). \tag{A31}$$

Last, the effect of excision is given by:

$$D_{ij}{}^{\mathrm{e}} \approx (1 - 2v)\, D_{ij}, \quad D_{i,j}{}^{\mathrm{e}} = 0. \tag{A32}$$

Equations A13 and A28–A32 now yield:

$$D_{ij} \approx \frac{\frac{u}{2L}\left(p_i + p_j + \sum_k D_{ik} + \sum_k D_{jk}\right) - \beta\, p_i p_j}{1 - (1 - r_{ij})\left[1 - 2\left(\alpha + \beta\overline{n}\right)\right]\left(1 - \frac{u}{2L}\,\overline{n}\right)(1 - 2v)}. \tag{A33}$$

Summing over all pairs of loci $i$ and $j$ and taking the limit as $L$ tends to infinity, the term $\frac{u}{2L}\,\overline{n}$ in the denominator vanishes. Furthermore, using $\overline{n} \approx (u - v - \alpha)/\beta$ so that $\alpha + \beta\overline{n} \approx u - v$, the denominator becomes approximately $r_{ij} + 2u$, giving:

$$2\sum_{i \neq j} D_{ij} \approx \mathcal{E}_1 \left[u\left(\overline{n} + 2\sum_{i \neq j} D_{ij}\right) - \beta\,\frac{\overline{n}^2}{2}\right] \tag{A34}$$

where $\mathcal{E}_1$ is the average over all possible pairs of insertion sites of $1/\left(r_{ij} + 2u\right)$. Again using $\overline{n} \approx (u - v - \alpha)/\beta$, we finally have:

$$2\sum_{i \neq j} D_{ij} \approx \frac{\mathcal{E}_1}{1 - u\,\mathcal{E}_1}\,\frac{u + v + \alpha}{2}\,\overline{n}\,. \tag{A35}$$

The inflation in variance caused by positive LD is thus given by:

$$\frac{\mathrm{Var}\,(n)}{\overline{n}} \approx 1 + \frac{\mathcal{E}_1}{1 - u\,\mathcal{E}_1}\,\frac{u + v + \alpha}{2}\,. \tag{A36}$$

Note that when linkage disequilibria make a significant contribution to the variance in $n$, a more accurate expression for the effect of selection on $\overline{n}$ can be obtained from equation A8:

$$\Delta_{\mathrm{sel}}\overline{n} \approx -\left(\alpha + \beta\overline{n}\right)\left(\overline{n} + 2\sum_{i \neq j} D_{ij}\right) \tag{A37}$$

which may be used together with equation A33 to obtain more precise (but complicated) expressions for $\overline{n}$ and $2\sum_{i \neq j} D_{ij}$.

In the case of a linear genetic map of length $R$ Morgans (with uniform density of recombination), $\mathcal{E}_1$ is given by:

$$\mathcal{E}_1 \approx \frac{2}{R^2} \int_0^R \frac{R-x}{x+2u}\, dx \tag{A38}$$

where the recombination rate $r$ is approximated by the genetic distance $x$, since $\mathcal{E}_1$ should be mostly generated by tightly linked loci as long as $R$ is not too large, yielding:

$$\mathcal{E}_1 \approx \frac{2}{R} \left[ (1+2\rho_R) \left[ \ln\left( \frac{1}{2} + \rho_R \right) - \ln(\rho_R) \right] - 1 \right] \tag{A39}$$

with $\rho_R = u/R$. When $R$ is large, the contribution of loosely linked loci to $\mathcal{E}_1$ becomes more important, and $x$ in the denominator of equation A38 must be replaced by $\left(1 - e^{-2x}\right)/2$ (Haldane, 1919), yielding a slightly more complicated expression (that tends to $1/\left(1/2 + 2u\right)$ as $R$ tends to infinity). Both expressions of $\mathcal{E}_1$ give similar results for values of $R$ up to 1, however, while the contribution of LD to the variance in the number of TEs per individual is generally very small when $R > 1$, as predicted by both expressions. Nevertheless, Figure S1 shows that replacing $x$ by $\left(1 - e^{-2x}\right)/2$ in the denominator of equation A38 yields more accurate predictions when $R$ is large.

**4. Partial clonality.** Assuming that at each generation, a proportion $\sigma$ of offspring is produced sexually (through random mating) while a proportion $1 - \sigma$ is produced clonally (through mitosis), the average number of TEs per genome at equilibrium stays approximately given by equation A12, while the effect of reproduction on pairwise genetic associations is now given by:

$$D_{ij}{}' = \left(1 - r_{ij}\,\sigma\right) D_{ij}{}^{\mathrm{s}} + r_{ij}\,\sigma\, D_{i,j}{}^{\mathrm{s}} \tag{A40}$$

$$D_{i,j}{}' = \left(1 - \sigma\right) D_{i,j}{}^{\mathrm{s}}\,. \tag{A41}$$

The effect of selection on $D_{ij}$ and $D_{i,j}$ is still given by equations A28 – A29, while the effects of transposition and excision are given by:

$$D_{ij}{}^{\mathrm{t}} \approx D_{ij}{}^{\mathrm{e}} + \frac{u}{2L} \left( p_i + p_j + \sum_k \left(D_{ik} + D_{i,k}\right) + \sum_k \left(D_{jk} + D_{j,k}\right) \right), \tag{A42}$$

$$D_{i,j}{}^{\mathrm{t}} \approx D_{i,j}{}^{\mathrm{e}} + \frac{u}{2L} \left( p_i + p_j + \sum_k \left(D_{ik} + D_{i,k}\right) + \sum_k \left(D_{jk} + D_{j,k}\right) \right), \tag{A43}$$

$$D_{ij}{}^{\mathrm{e}} \approx (1-2v)\, D_{ij}, \quad D_{i,j}{}^{\mathrm{e}} \approx (1-2v)\, D_{i,j}. \tag{A44}$$

Assuming that the rate of sex $\sigma$ is small, one obtains, at equilibrium:

$$\begin{aligned} D_{ij} + D_{i,j} \approx\ & \frac{4u + \sigma\,(1+2r_{ij})}{(2u+\sigma)\,(2u + r_{ij}\,\sigma)} \\ & \times \left[ \frac{u}{2L} \left( p_i + p_j + \sum_k (D_{ik} + D_{i,k}) + \sum_k (D_{jk} + D_{j,k}) \right) - \beta\, p_i p_j \right] \end{aligned} \tag{A45}$$

and the contribution of genetic associations to the variance in $n$ is thus given by:

$$2 \sum_{i \neq j} (D_{ij} + D_{i,j}) \approx \frac{\mathcal{E}_2}{1 - u\,\mathcal{E}_2} \frac{u+v+\alpha}{2}\, \overline{n} \tag{A46}$$

where $\mathcal{E}_2$ is the average over all pairs of possible insertion sites of the fraction on the first line of equation A45. When the rate of sex $\sigma$ is small, distant pairs of loci may contribute significantly to the average, which is thus computed as:

$$\mathcal{E}_2 = \frac{2}{R^2} \int_0^R \frac{(R-x)\,[4u + \sigma\,[1 + 2r(x)]]}{(2u+\sigma)\,[2u + r(x)\,\sigma]}\, dx \tag{A47}$$

in the case of a linear genetic map of length $R$, where $r(x)$ is given by Haldane's mapping function (Haldane, 1919): $r(x) = [1 - e^{-2x}]\,/2$. Equation A47 yields:

$$\mathcal{E}_2 = \frac{2R\,[R\,(1+4\rho_\sigma) - \ln(-4\rho_\sigma)] + \mathrm{Li}_2(1+4\rho_\sigma) - \mathrm{Li}_2\big(e^{2R}\,(1+4\rho_\sigma)\big)}{R^2\sigma\,(1+2\rho_\sigma)\,(1+4\rho_\sigma)} \tag{A48}$$

where $\rho_\sigma = u/\sigma$, and $\mathrm{Li}_2(x)$ is the polylogarithm function $\sum_{k=1}^{\infty} z^k/k^2$. The inflation in variance caused by positive genetic associations between loci is thus given by:

$$\frac{\mathrm{Var}\,(n)}{\overline{n}} \approx 1 + \frac{\mathcal{E}_2}{1 - u\,\mathcal{E}_2} \frac{u+v+\alpha}{2}\,. \tag{A49}$$

**5. Partial selfing.** Assuming that a proportion $s$ of offspring is produced by selfing, while a proportion $1-s$ is produced by random mating, the change in frequency of insertions at site $i$ is now given by (from equations A6 and A7):

$$\Delta_{\mathrm{sel}} p_i \approx -\,(\alpha + \beta\overline{n})\,(p_i q_i + D_{i,i})\,. \tag{A50}$$

The association $D_{i,i}$ measures the excess homozygosity at site $i$ caused by selfing, and equals $F p_i q_i$ to leading order, where $F = s/\,(2-s)$ is the inbreeding coefficient. Therefore,

$$\Delta_{\mathrm{sel}} \overline{n} \approx -\,(\alpha + \beta\overline{n})\,(1+F)\,\overline{n}, \tag{A51}$$

while the change in $\overline{n}$ due to transposition and excision is still given by $(u - v)\,\overline{n}$, giving, at equilibrium:

$$\overline{n} \approx \frac{u - v - \alpha\,(1 + F)}{\beta\,(1 + F)}. \tag{A52}$$

The coefficient $a_i = \alpha + \beta\overline{n}$ of equation A6 is thus now approximately $(u - v)\,/\,(1 + F)$ at equilibrium, while the net effect of selection against each element, $(\alpha + \beta\overline{n})\,(1 + F)$ (from equation A50) stays approximately equal to $u - v$. The effect of reproduction on pairwise genetic associations is given by:

$$D_{ij}{}' = (1 - r_{ij})\,D_{ij}{}^{\mathrm{s}} + r_{ij}\,D_{i,j}{}^{\mathrm{s}} \tag{A53}$$

$$D_{i,j}{}' = \frac{s}{2}\left(D_{ij}{}^{\mathrm{s}} + D_{i,j}{}^{\mathrm{s}}\right). \tag{A54}$$

From equations A6 and A14, and neglecting associations involving more than two loci, one obtains:

$$\begin{aligned} D_{ij}{}^{\mathrm{s}} \approx {} & [1 - 2\,(\alpha + \beta\overline{n})]\,D_{ij}{}^{\mathrm{t}} - (\alpha + \beta\overline{n})\left(D_{ij,i}{}^{\mathrm{t}} + D_{ij,j}{}^{\mathrm{t}}\right) \\ & - \beta\left(D_{iijj}{}^{\mathrm{t}} + D_{ij,ij}{}^{\mathrm{t}} + D_{iij,j}{}^{\mathrm{t}} + D_{ijj,i}{}^{\mathrm{t}}\right). \end{aligned} \tag{A55}$$

The associations appearing on the second line of equation A55 are given by $D_{iijj}{}^{\mathrm{t}} \approx p_i p_j$, $D_{iij,j}{}^{\mathrm{t}} = D_{ijj,i}{}^{\mathrm{t}} \approx F p_i p_j$ and $D_{ij,ij}{}^{\mathrm{t}} \approx \phi_{ij} p_i p_j$, where $\phi_{ij}$ is the probability of joint identity-by-descent at loci $i$ and $j$ (that equals 1 under complete selfing). In the following, we will consider that the outcrossing rate $o = 1 - s$ is small, in which case one can show that $D_{ij,i}{}^{\mathrm{t}} \approx F\,D_{ij}{}^{\mathrm{t}}$, leading to:

$$D_{ij}{}^{\mathrm{s}} \approx [1 - 2\,(\alpha + \beta\overline{n})\,(1 + F)]\,D_{ij}{}^{\mathrm{t}} - \beta\,(1 + 2F + \phi_{jk})\,p_j p_k. \tag{A56}$$

Similarly, we have:

$$D_{i,j}{}^{\mathrm{s}} \approx [1 - 2\,(\alpha + \beta\overline{n})]\,D_{i,j}{}^{\mathrm{t}} + 2F\,(\alpha + \beta\overline{n})\,D_{ij}{}^{\mathrm{t}} - \beta\,(1 + 2F + \phi_{jk})\,p_j p_k. \tag{A57}$$

The effects of transposition and excision are given by:

$$D_{ij}{}^{\mathrm{t}} \approx D_{ij}{}^{\mathrm{e}} + \frac{u}{2L}\left[(p_i + p_j)\,(1 + F) + \sum_k (D_{ik} + D_{i,k}) + \sum_k (D_{jk} + D_{j,k})\right], \tag{A58}$$

$$D_{i,j}{}^{\mathrm{t}} \approx D_{i,j}{}^{\mathrm{e}} + \frac{u}{2L}\left[(p_i + p_j)\,(1 + F) + \sum_k (D_{ik} + D_{i,k}) + \sum_k (D_{jk} + D_{j,k})\right], \tag{A59}$$

$$D_{ij}{}^{\mathrm{e}} \approx (1 - 2v)\,D_{ij}, \quad D_{i,j}{}^{\mathrm{e}} \approx (1 - 2v)\,D_{i,j}. \tag{A60}$$

Assuming that the outcrossing rate $o$ is small (so that $\phi_{ij} \approx F \approx 1 - 2o$), one obtains, at equilibrium:

$$D_{ij} + D_{i,j} \approx \frac{1 + 2r_{ij}}{o r_{ij} + u\,(1 + 2r_{ij})} \times \left[ \frac{u}{2L} \left( 2p_i + 2p_j + \sum_k (D_{ik} + D_{i,k}) + \sum_k (D_{jk} + D_{j,k}) \right) - 4\beta\, p_i p_j \right] \tag{A61}$$

and the contribution of genetic associations between TEs at different sites to the variance in $n$ is thus given by:

$$2 \sum_{i \neq j} (D_{ij} + D_{i,j}) \approx \frac{\mathcal{E}_3}{1 - u\,\mathcal{E}_3} (u + v + 2\alpha)\, \overline{n} \tag{A62}$$

where $\mathcal{E}_3$ is the average over all pairs of possible insertion sites of the fraction on the first line of equation A61. In the case of a linear map of length $R$, we have:

$$\mathcal{E}_3 = \frac{2}{R^2} \int_0^R \frac{(R - x)\,[1 + 2r(x)]}{o r(x) + u\,[1 + 2r(x)]}\, dx \tag{A63}$$

with $r(x) = \left[1 - e^{-2x}\right]/2$, yielding:

$$\mathcal{E}_3 = \frac{2R \left[ R\,(1 + 4\rho_o) - \ln\!\left( -\frac{2\rho_o}{1+2\rho_o} \right) \right] + \mathrm{Li}_2\!\left( \frac{1+4\rho_o}{1+2\rho_o} \right) - \mathrm{Li}_2\!\left( e^{2R} \frac{1+4\rho_o}{1+2\rho_o} \right)}{R^2 o\,(1 + 2\rho_o)\,(1 + 4\rho_o)} \tag{A64}$$

where $\rho_o = u/o$ and $\mathrm{Li}_2(x)$ is the polylogarithm function $\sum_{k=1}^{\infty} z^k/k^2$. Under weak outcrossing, the inflation in variance caused by genetic associations between loci is thus given by:

$$\frac{\mathrm{Var}\,(n)}{\overline{n}\,(1 + F)} \approx 1 + \frac{\mathcal{E}_3}{1 - u\,\mathcal{E}_3} \, \frac{u + v + 2\alpha}{2}\,. \tag{A65}$$

**6. The Hill-Robertson effect.** In a finite population, the Hill-Robertson effect tends to generate negative LD between TE copies. Assuming that the mean number of TEs per genome $\overline{n}$ is sufficiently large, so that the coefficient of epistasis $\beta$ is small relative to the coefficient of selection $a_i = \alpha + \beta\overline{n}$, the contribution of the Hill-Robertson effect to the linkage disequilibrium between TE copies should be approximately the same as between two deleterious alleles with additive fitness effects ($h = 1/2$), replacing $-sh$ by $a_i$. Denoting $\langle X \rangle$ the expected value of quantity $X$ at mutation–selection–drift equilibrium and assuming that deleterious alleles stay near mutation–selection balance (at low frequency), Roze (2021) showed that an approximation for the average LD $\langle D_{ij} \rangle$ generated by the Hill-Robertson effect between two deleterious alleles with selection coefficients $s_i$, $s_j$ and dominance coefficients $h_i$, $h_j$ in a population of size $N$ is given by:

$$\left\langle {D_{ij}}^2 \right\rangle \approx \frac{p_i p_j}{4N \left(r_{ij} + s_i h_i + s_j h_j\right)} \tag{A66}$$

$$\langle p_i D_{ij} \rangle \approx -\frac{s_j h_j \left\langle {D_{ij}}^2 \right\rangle}{r_{ij} + 2 s_i h_i + s_j h_j} \tag{A67}$$

$$\langle D_{ij} \rangle \approx \frac{s_i \left(2h_i - d_i\right) \langle p_i D_{ij} \rangle + s_j \left(2h_j - d_j\right) \langle p_j D_{ij} \rangle}{r_{ij} + s_i h_i + s_j h_j} \tag{A68}$$

where $d_i = 1 - 2h_i$. These equations can be adapted to our model of TE dynamics by setting $d_i$, $d_j$ to zero (no dominance), and replacing $s_i h_i$ and $s_j h_j$ in the numerators of equations A67 – A68 by $\alpha + \beta \overline{n} \approx u - v$. However, because excision tends to reduce insertion frequencies and associations, extra terms in $v$ appear in the denominators, so that $s_i h_i$ and $s_j h_j$ in the denominators of equations A66 – A68 must simply be replaced by $u$. Combining the different sources of LD (transposition, epistasis, Hill-Robertson effect) and assuming that the genome map length is sufficiently large so that $\sum_{i \neq j} D_{ij}$ remains small, one obtains:

$$\langle D_{ij} \rangle \approx \frac{\frac{u}{2L} \left(p_i + p_j\right) - \beta\, p_i p_j}{r_{ij} + 2u} - \frac{\left(u - v\right)^2 p_i p_j}{N \left(r_{ij} + 2u\right)^2 \left(r_{ij} + 3u\right)}. \tag{A69}$$

Approximating $p_i$ and $p_j$ by the frequencies of insertions at sites $i$ and $j$ in the deterministic limit, $\left(u - v - \alpha\right) / \left(2L\beta\right)$, yields:

$$\frac{\langle D_{ij} \rangle}{\langle pq_{ij} \rangle} \approx \frac{\left(u + v + \alpha\right) \beta}{\left(u - v - \alpha\right) \left(r_{ij} + 2u\right)} - \frac{\left(u - v\right)^2}{N \left(r_{ij} + 2u\right)^2 \left(r_{ij} + 3u\right)} \tag{A70}$$

with $pq_{ij} = p_i q_i p_j q_j$. This result can be extended to the case of partially selfing populations and tightly linked loci using separation of timescales arguments (e.g., Nordborg, 1997; Roze, 2016; Stetsenko and Roze, 2022): this is achieved by replacing $N$ by $N / \left(1 + F\right)$ and $r_{ij}$ by $r_{ij} \left(1 - F\right)$ in the right part of equation A70 (note that the effective strength of selection against TE insertions is not affected by selfing and stays $\approx u - v$, as shown above). From this, and using the expressions given in the previous subsection for deterministic effects, one obtains:

$$\begin{aligned} \frac{\langle D_{ij} \rangle}{\langle pq_{ij} \rangle} \approx & \frac{\left[\left[1 + F \left(1 + 2F\right)\right] u + \left(1 + 3F\right) \left[v + \alpha \left(1 + F\right)\right]\right] \beta}{\left[u - v - \alpha \left(1 + F\right)\right] \left[r_{ij} \left(1 - F\right) + 2u\right]} \\ & - \frac{\left(u - v\right)^2 \left(1 + F\right)}{N \left[r_{ij} \left(1 - F\right) + 2u\right]^2 \left[r_{ij} \left(1 - F\right) + 3u\right]} \end{aligned} \tag{A71}$$

which can be expressed in terms of the outcrossing rate $o$ using $F = (1-o)/(1+o)$.

**7. Hill-Robertson effect: more accurate expression for small map length.** When deleterious alleles segregate at many linked loci, $N$ in equation A66 should be replaced by the effective population size $N_\mathrm{e}$, which is reduced by background selection (Charlesworth et al., 1993; Charlesworth, 1996). When recombination is sufficiently high, and assuming a uniform rate of mutation and crossing-over along the chromosome, $N_\mathrm{e}$ should be approximately the same at all loci (e.g., Roze, 2021). In the case of a tight linkage map, however, $N_\mathrm{e}$ may be different at loci $i$ and $j$ (depending on their position along the chromosome). In order to incorporate the effect of background selection on $\langle D_{ij} \rangle$ in this situation, we can extend the method used to compute the effect of background selection on diversity at a neutral locus (e.g., Hudson and Kaplan, 1995), namely, compute the effect of a third selected locus $k$ on $\langle D_{ij} \rangle$, and assume that all other loci have multiplicative effects of $\langle D_{ij} \rangle$. Assuming random mating and measuring genetic associations at the gametic stage, the method of Roze (2021) can be extended to include a third selected locus, yielding (see *Mathematica* notebook available as Supplementary Material for derivation):

$$\left\langle {D_{ijk}}^2 \right\rangle \approx \frac{p_i p_j p_k}{4N\left(r_{ijk} + s_i h_i + s_j h_j + s_k h_k\right)} \tag{A72}$$

$$\left\langle D_{ij} D_{ijk} \right\rangle \approx -\frac{s_k h_k \left\langle {D_{ijk}}^2 \right\rangle}{r_{ijk} + r_{ij} + 2 s_i h_i + 2 s_j h_j + s_k h_k} \tag{A73}$$

$$\left\langle D_{ij} D_{ik} \right\rangle \approx -\frac{s_j h_j \left\langle D_{ij} D_{ijk} \right\rangle + s_k h_k \left\langle D_{ik} D_{ijk} \right\rangle}{r_{ij} + r_{ik} + 2 s_i h_i + s_j h_j + s_k h_k} \tag{A74}$$

$$\left\langle p_i D_{ijk} \right\rangle \approx -\frac{s_j h_j \left\langle D_{ij} D_{ijk} \right\rangle + s_k h_k \left\langle D_{ik} D_{ijk} \right\rangle}{r_{ijk} + 2 s_i h_i + s_j h_j + s_k h_k} \tag{A75}$$

$$\begin{aligned}\left\langle D_{ijk} \right\rangle \approx\ & \frac{1}{r_{ijk} + s_i h_i + s_j h_j + s_k h_k} \\ & \times \Big[ 2 s_i h_i \left\langle D_{ij} D_{ik} \right\rangle + 2 s_j h_j \left\langle D_{ij} D_{jk} \right\rangle + 2 s_k h_k \left\langle D_{ik} D_{jk} \right\rangle \\ & + s_i \left(2h_i - d_i\right) \left\langle p_i D_{ijk} \right\rangle + s_j \left(2h_j - d_j\right) \left\langle p_j D_{ijk} \right\rangle \\ & + s_k \left(2h_k - d_k\right) \left\langle p_k D_{ijk} \right\rangle \Big]\end{aligned} \tag{A76}$$

$$\left\langle {D_{ij}}^2 \right\rangle \approx \frac{p_i p_j}{4N\left(r_{ij} + s_i h_i + s_j h_j\right)} - \frac{s_k h_k \left\langle D_{ij} D_{ijk} \right\rangle}{r_{ij} + s_i h_i + s_j h_j} \tag{A77}$$

$$\left\langle p_i D_{ij} \right\rangle \approx -\frac{s_j h_j \left\langle {D_{ij}}^2 \right\rangle + s_k h_k \left(\left\langle D_{ij} D_{ik} \right\rangle + \left\langle p_i D_{ijk} \right\rangle\right)}{r_{ij} + 2 s_i h_i + s_j h_j} \tag{A78}$$

$$\langle D_{ij} \rangle \approx \frac{1}{r_{ij} + s_i h_i + s_j h_j} \times \Big[ s_i (2h_i - d_i) \langle p_i D_{ij} \rangle + s_j (2h_j - d_j) \langle p_j D_{ij} \rangle - s_k h_k \langle D_{ijk} \rangle - s_k d_k \langle p_k D_{ijk} \rangle \Big] \tag{A79}$$

where $r_{ijk} = (r_{ij} + r_{ik} + r_{jk})/2$ is the probability that at least one recombination event occurs between loci $i$, $j$ and $k$ (note that equations A72 – A79 remain valid for any ordering of the three loci along the chromosome). As before, these equations can be adapted to our model of TE dynamics by setting $d_i$, $d_j$ and $d_k$ to zero (no dominance), replacing $s_i h_i$, $s_j h_j$ and $s_k h_k$ in the numerators of equations A72 – A79 by $\alpha + \beta \overline{n} \approx u - v$, and replacing $s_i h_i$, $s_j h_j$ and $s_k h_k$ in the denominators by $u$. One obtains:

$$\langle D_{ij} \rangle \approx -\frac{(u-v)^2 p_i p_j}{N (r_{ij} + 2u)^2 (r_{ij} + 3u)} \left[ 1 + (u-v)^2 T_{ijk} p_k \right] \tag{A80}$$

where $T_{ijk}$ is a positive function of $r_{ij}$, $r_{ik}$, $r_{jk}$ and $u$ (available in the *Mathematica* notebook available as Supplementary Material). Assuming that all TEs at different sites have multiplicative effects on $\langle D_{ij} \rangle$ yields:

$$\langle D_{ij} \rangle \approx -\frac{(u-v)^2 p_i p_j}{N (r_{ij} + 2u)^2 (r_{ij} + 3u)} \exp\left[ \frac{(u-v)^2 \overline{n}}{2R} \int_0^R T_{ijk} \, dx_k \right] \tag{A81}$$

where $x_k$ is the position of insertion site $k$ along the genetic map (from 0 to $R$). Combining equations A33 and A81 in order to take into account the joint effects of transposition, epistasis and the Hill-Robertson effect yields:

$$\langle D_{ij} \rangle \approx \frac{\frac{u}{2L} \left( p_i + p_j + \sum_k \langle D_{ik} \rangle + \sum_k \langle D_{jk} \rangle \right) - \beta \, p_i p_j}{r_{ij} + 2u} - \frac{(u-v)^2 p_i p_j}{N (r_{ij} + 2u)^2 (r_{ij} + 3u)} \exp\left[ \frac{(u-v)^2 \overline{n}}{2R} \int_0^R T_{ijk} \, dx_k \right], \tag{A82}$$

giving:

$$2 \sum_{i \neq j} D_{ij} \approx \frac{1}{1 - u \, \mathcal{E}_1} \left[ \mathcal{E}_1 \frac{u + v + \alpha}{2} \overline{n} - \mathcal{E}_4 \frac{(u-v)^2}{2N} \overline{n}^2 \right], \tag{A83}$$

where $\mathcal{E}_1$ is given by equation A39, while

$$\mathcal{E}_4 = \frac{1}{R^2} \iint_0^R \frac{\exp\left[ \frac{(u-v)^2 \overline{n}}{2R} \int_0^R T_{ijk} \, dx_k \right]}{(r_{ij} + 2u)^2 (r_{ij} + 3u)} \, dx_i \, dx_j \tag{A84}$$

which can be computed numerically using *Mathematica* (see Supplementary Material). For this, $r_{ij}$, $r_{ik}$ and $r_{jk}$ are approximated by $|x_j - x_i|$, $|x_k - x_i|$ and $|x_k - x_j|$, respectively. From equation A83, the relative effect of LD on the variance in $n$ is given by:

$$\frac{\mathrm{Var}(n)}{\overline{n}} \approx \frac{1}{1 - u\,\mathcal{E}_1} \left[ \mathcal{E}_1 \frac{u + v + \alpha}{2} - \mathcal{E}_4 \frac{(u-v)^2}{2N} \frac{u - v - \alpha}{\beta} \right]. \tag{A85}$$

One can note that equations A83 and A85 only depend on the $Nu$, $Nv$, $N\alpha$, $N\beta$ and $NR$ products. In particular, equation A85 can be written as:

$$\frac{\mathrm{Var}(n)}{\overline{n}} \approx \frac{1}{1 - Nu\,\tilde{\mathcal{E}}_1} \left[ \tilde{\mathcal{E}}_1 \frac{Nu + Nv + N\alpha}{2} - \tilde{\mathcal{E}}_4 \frac{(Nu - Nv)^2 (Nu - Nv - N\alpha)}{2N\beta} \right] \tag{A86}$$

with:

$$\tilde{\mathcal{E}}_1 = \frac{\mathcal{E}_1}{N} = \frac{2}{(NR)^2} \int_0^{NR} \frac{NR - y}{y + 2Nu}\, dy, \tag{A87}$$

$$\tilde{\mathcal{E}}_4 = \frac{\mathcal{E}_4}{N^3} = \frac{1}{(NR)^2} \iint_0^{NR} \frac{\exp\left[\frac{(Nu-Nv)^2\overline{n}}{2NR} \int_0^{NR} \tilde{T}_{ijk}\, dy_k\right]}{(Nr_{ij} + 2Nu)^2 (Nr_{ij} + 3Nu)}\, dy_i\, dy_j, \tag{A88}$$

$\tilde{T}_{ijk} = T_{ijk}/N^2$ and $Nr_{ij} = |y_j - y_i|$.

**8. Extension to multiple TE families.** The previous analyses can be extended to the case where $f$ different TE families co-exist in the genome. Assuming for simplicity that $\alpha$, $\beta$, $u$ and $v$ are the same for all families and that ectopic recombination can only occur between TEs from the same family, the fitness of an individual is given by:

$$W = \exp\left( -\alpha \sum_{y=1}^{f} n_y - \beta \sum_{y=1}^{f} n_{\mathrm{p},y} \right) \tag{A89}$$

where $n_y$ is the number of TEs from family $y$, and $n_{\mathrm{p},y}$ the number of pairs of TEs from family $y$ present in the genome. Denoting $D_{i_y j_z}$ the linkage disequilibrium between TEs from families $y$ and $z$ present at sites $i$ and $j$, and using the results above, the sum of all LD between transposons from the same family is given by:

$$2 \sum_y \sum_{i \neq j} D_{i_y j_y} \approx \frac{f}{1 - u\,\mathcal{E}_1} \left[ \mathcal{E}_1 \frac{u + v + \alpha}{2} \overline{n} - \mathcal{E}_4 \frac{(u-v)^2}{2N} \overline{n}^2 \right] \tag{A90}$$

where $\mathcal{E}_4$ is now given by:

$$\mathcal{E}_4 = \frac{1}{R^2} \iint_0^{R} \frac{\exp\left[\frac{(u-v)^2 \overline{n} f}{2R} \int_0^{R} T_{ijk}\, dx_k\right]}{(r_{ij} + 2u)^2 (r_{ij} + 3u)}\, dx_i\, dx_j\,. \tag{A91}$$

The sum of LD between transposons from different families is only generated by the Hill-Robertson effect, and thus given by:

$$2\sum_{y\neq z}\sum_{i\neq j} D_{i_y j_z} \approx -f\,(f-1)\,\frac{\mathcal{E}_4}{1-u\,\mathcal{E}_1}\,\frac{(u-v)^2}{2N}\,\overline{n}^2\,. \tag{A92}$$

**9. Evolution of recombination: deterministic model.** We now consider how transposable elements present along a linear chromosome will affect the evolution of a modifier affecting the genetic map length $R$ of the chromosome, in an infinite population. We assume that two alleles $M$ and $m$ segregate at the modifier locus, allele $m$ increasing the chromosome map length by an amount $\delta R/2$ when heterozygous and $\delta R$ when homozygous. The following derivations assume that $R$ stays sufficiently large, so that the contribution of linkage disequilibria to the variance in TE number among individuals remains small (e.g., $NR \geq 10^3$ for the parameter values used in Figure 1). For simplicity, the modifier is supposed to be located at the mid-point of the chromosome (but the exact position of the modifier should not affect too much the results as long as $R$ is not too small). From equations 1 and 10 in the main text, the fitness of an individual can be written as:

$$\begin{aligned} W = \exp\Bigg[&-\alpha\sum_{i}\left(X_{i,\varnothing}+X_{\varnothing,i}\right) \\ &-\frac{\tilde{\beta}}{2}\left[1-\theta+\theta\left[R+\frac{\delta R}{2}\left(X_{m,\varnothing}+X_{\varnothing,m}\right)\right]\right]\sum_{i\neq j}\left(X_{i,\varnothing}+X_{\varnothing,i}\right)\left(X_{j,\varnothing}+X_{\varnothing,j}\right)\Bigg] \end{aligned} \tag{A93}$$

where $X_{m,\varnothing}$ and $X_{\varnothing,m}$ equal 1 if allele $m$ is present on the first or second haplotype of the individual, and 0 otherwise. To the first order in $\epsilon$, we thus have:

$$
\begin{aligned}
\frac{W}{\overline{W}} = 1 &- \left(\alpha + \overline{\beta}\,\overline{n}\right) \sum_i \left(\zeta_{i,\varnothing} + \zeta_{\varnothing,i}\right) \\
&- \frac{\overline{\beta}}{2} \sum_{i \neq j} \left(\zeta_{ij,\varnothing} + \zeta_{\varnothing,ij} - 2D_{ij} + \zeta_{i,j} + \zeta_{j,i} - 2D_{i,j}\right) \\
&- \frac{\delta R}{2} \frac{\tilde{\beta}\theta \overline{n}^2}{2} \left(\zeta_{m,\varnothing} + \zeta_{\varnothing,m}\right) \\
&- \frac{\delta R}{2} \tilde{\beta}\theta \overline{n} \sum_i \left(\zeta_{mi,\varnothing} + \zeta_{\varnothing,mi} - 2D_{mi} + \zeta_{m,i} + \zeta_{i,m} - 2D_{m,i}\right) \\
&- \frac{\delta R}{2} \frac{\tilde{\beta}\theta}{2} \sum_{i \neq j} \Big(\zeta_{mij,\varnothing} + \zeta_{\varnothing,mij} - 2D_{mij} + \zeta_{mi,j} + \zeta_{j,mi} - 2D_{mi,j} \\
&\qquad + \zeta_{mj,i} + \zeta_{i,mj} - 2D_{mj,i} + \zeta_{m,ij} + \zeta_{ij,m} - 2D_{m,ij}\Big) + o(\epsilon)
\end{aligned}
\tag{A94}
$$

with $\overline{\beta} = \beta + \delta R \tilde{\beta} \theta p_m$. The change in frequency of allele $m$ over a generation is given by:

$$
\Delta p_m = \mathrm{E}\left[\frac{W}{\overline{W}} \frac{\zeta_{m,\varnothing} + \zeta_{\varnothing,m}}{2}\right]. \tag{A95}
$$

Using A94 and assuming that $\sum_{i \neq j} D_{ij}$ stays small relative to $\overline{n}^2$, this yields, to the first order in $\delta R$:

$$
\Delta p_m \approx -\frac{\delta R}{2} \frac{\tilde{\beta}\theta \overline{n}^2}{2} p_m q_m - (\alpha + \beta n) \sum_i D_{mi} - \frac{\beta}{2} \sum_{i \neq j} D_{mij}\,. \tag{A96}
$$

If an inherent fitness cost $c$ per crossover is included into the model, an extra term $-\left(\delta R/2\right) c p_m q_m$ must be added to equation A96. The first term of equation A96 corresponds to the effect of the direct fitness cost of recombination generated by ectopic recombination events among TEs. The second term favors the modifier allele that tends to be less associated with TE insertions (i.e., that tends to be found on better purged genetic backgrounds), while the third term favors the modifier allele that tends to be more associated with intermediate genotypes (over all possible pairs of polymorphic insertion sites) than with extreme genotypes, given that intermediate genotypes have a higher mean fitness than extreme genotypes under negative epistasis.

Throughout the following, recombination rates are assumed to be small, as indirect selection on the modifier should mostly be driven by tightly linked loci (Barton, 1995; Roze, 2021). In this case, a recurrence equation on $D_{mi}$ is given by:

$$
D_{mi}' \approx \left(1 - r_{mi}\right) D_{mi}^{\mathrm{s}} \tag{A97}
$$

with

$$D_{mi}{}^{\mathrm{s}} \approx \mathrm{E}\left[\frac{W}{\overline{\overline{W}}}\frac{\zeta_{mi,\varnothing}+\zeta_{\varnothing,mi}}{2}\right]. \tag{A98}$$

From equation A94, using $\alpha+\beta\overline{n} \approx u-v$ and assuming $\overline{n} \gg 1$ (so that $\beta\overline{n} \gg \beta$), one obtains:

$$D_{mi}{}^{\mathrm{s}} \approx [1-(u-v)]\,D_{mi}{}^{\mathrm{t}} - (u-v)\sum_{j\neq i} D_{mij} - \frac{\delta R}{2}\tilde{\beta}\theta\overline{n}p_i p_m q_m \tag{A99}$$

(the term in $\delta R$ being generated from the term on the fourth line of equation A94). The effect of transposition on $D_{mi}$ is found to be negligible under our assumptions, while excision multiplies $D_{mi}$ by a factor $1-v$, yielding at equilibrium:

$$D_{mi} \approx -\frac{u-v}{r_{mi}+u}\sum_{j\neq i} D_{mij} - \frac{\delta R}{2}\frac{\tilde{\beta}\theta\overline{n}}{r_{mi}+u}p_i p_m q_m\,. \tag{A100}$$

The first term of equation A100 shows that allele $m$ tends to be found on better purged backgrounds ($D_{mi}<0$) if $m$ is more often associated than $M$ with extreme genotypes at pairs of segregating insertions sites ($D_{mij}>0$), as selection is more efficient among extreme genotypes. Additionally, $m$ tends to be found on better purged backgrounds if it increases selection against TEs by increasing the rate of ectopic recombination (i.e., if $\theta>0$; second term of equation A100).

Under the same assumptions, a recurrence equation on $D_{mij}$ is given by (e.g., Barton, 1995):

$$D_{mij}{}' \approx (1-r_{mij})\,D_{mij}{}^{\mathrm{s}} - \frac{\delta r_{ij}}{2}D_{ij}\,p_m q_m \tag{A101}$$

where $r_{mij} = (r_{mi}+r_{mj}+r_{ij})/2$ is the probability that at least one recombination event occurs among the three loci, and $\delta r_{ij}$ the effect of allele $m$ on $r_{ij}$. We then have (Barton and Turelli, 1991):

$$\begin{aligned} D_{mij}{}^{\mathrm{s}} &\approx \mathrm{E}\left[\frac{W}{\overline{\overline{W}}}\frac{\zeta_{mij,\varnothing}+\zeta_{\varnothing,mij}}{2}\right] - \Delta p_m D_{ij} \\ &\approx \mathrm{E}\left[\frac{W}{\overline{\overline{W}}}\frac{\zeta_{mij,\varnothing}+\zeta_{\varnothing,mij}}{2}\right] + \frac{\delta R}{2}\frac{\tilde{\beta}\theta\overline{n}^2}{2}D_{ij}\,p_m q_m\,. \end{aligned} \tag{A102}$$

Using equation A94, one arrives at:

$$D_{mij}{}^{\mathrm{s}} \approx [1-2(u-v)]\,D_{mij}{}^{\mathrm{t}} - \delta R\tilde{\beta}\theta\overline{n}D_{ij}\,p_m q_m - \frac{\delta R}{2}\tilde{\beta}\theta p_i p_j p_m q_m, \tag{A103}$$

the second and third terms being generated by the fourth and fifth lines of equation A94, respectively. Again, the effect of transposition on $D_{mij}$ can be neglected under our assumptions, while excision multiplies $D_{mij}$ by a factor $(1-v)^2 \approx 1-2v$, giving at equilibrium:

$$D_{mij} \approx -\frac{1}{r_{mij}+2u}\left(\frac{\delta r_{ij}}{2}D_{ij} + \delta R\tilde{\beta}\theta\overline{n}D_{ij} + \frac{\delta R}{2}\tilde{\beta}\theta p_i p_j\right)p_m q_m. \qquad \text{(A104)}$$

The first term within the brackets of equation A104 corresponds to the fact that LD among insertion sites tend to be weaker in the background of allele $m$, as this allele increases recombination. The fact that $m$ increases the rate of ectopic recombination and therefore the strength of selection against TEs (when $\theta > 0$) also reduces LD in the background of allele $m$, by decreasing the frequency of insertions (second term). Finally, the third term corresponds to the fact that by increasing ectopic recombination, $m$ increases the strength of negative epistasis among insertions, which tends to make LD less positive in the $m$ than in the $M$ background.

Using the fact that recombination rates can be approximated by genetic distances in the case of closely linked loci, we have $\delta r_{ij} \approx \delta R\, r_{ij}/R$ (Roze, 2021), while under our assumptions the linkage disequilibrium $D_{ij}$ is approximately:

$$D_{ij} \approx \frac{1}{r_{ij}+2u}\left[\frac{u}{2L}(p_i+p_j) - \beta\, p_i p_j\right]. \qquad \text{(A105)}$$

Using $2\sum_i p_i = \overline{n}$, equations A96, A100, A104 and A105 yield an expression for the change in frequency of the modifier in terms of the different parameters of the model and of various averages of functions of recombination rates, that can be obtained by numerical integration over the genetic map (see *Mathematica* notebook available as Supplementary Material).

When $\theta = 0$ (no effect of the modifier on the rate of ectopic recombination), the results take a similar form as in Barton's (1995) model on the effect of epistasis on selection for recombination, except that in the present model $D_{ij}$ is positive despite the fact that epistasis is negative. As a consequence, the frequency of extreme genotypes at pairs of insertions sites is lower in the $m$ than in the $M$ background ($D_{mij} < 0$), which benefits the modifier since extreme genotypes have a lower average fitness than intermediate genotypes under negative epistasis (third term of equation A96) — increasing recombination thus tends to increase the mean fitness of offspring. However, reducing the frequency of extreme genotypes decreases the efficiency of selection against TEs (increasing recombination decreases the variance in fitness among offspring), causing a higher TE load on the $m$ background (first term of equation A100), which disfavors allele $m$ (second term of equation A96). Overall, the second effect is generally stronger than the first, so that indirect selection disfavors recombination due to the higher TE load associated with $m$. Indirect selection may become positive for some parameter values (the effect on mean fitness being stronger than the effect on the variance in fitness), but only when $R$ is large and $\overline{n}$ is small, in which case indirect selection is extremely weak and easily overwhelmed by any slight direct fitness effect of the modifier. In parameter regions where indirect selection is stronger (lower $R$ and/or higher $\overline{n}$), it tends to be dominated by the effect of recombination on the variance in fitness among offspring (second term of equation A96), which from the above equations takes the form $s_{\text{ind,det}}\, p_m q_m$ with:

$$s_{\text{ind,det}} \approx -\frac{\delta R}{2R} (u-v)^2 \, \mathcal{E}\left[\frac{r_{ij}}{(r_{mi}+u)(r_{mij}+2u)(r_{ij}+2u)}\right] \frac{u+v+\alpha}{4} \overline{n} \qquad \text{(A106)}$$

where $\mathcal{E}$ is the average over all possible pairs of insertions sites $i$ and $j$.

When $\theta > 0$, allele $m$ has the additional effect of increasing the rate of ectopic recombination. This generates a direct selective pressure against $m$ due to the fitness cost of ectopic recombination (first term of equation A96), but also several indirect effects: (i) TEs are purged more efficiently (second term of equation A100), (ii) LD among TEs is reduced due to their elimination (second term of equation A104), (iii) LD among TEs tends to be less positive in the $m$ background due to the stronger negative epistasis (third term of equation A104). Effect (i) favors allele $m$, while effects (ii) and (iii) favor $m$ through the effect on the mean fitness of offspring (third term of equation A96), but disfavor $m$ through the effect on the variance in fitness among offspring (second term of equation A96). Numerical analysis show that these indirect effects are always weaker than the direct fitness cost of ectopic recombination, however (while among indirect effects, (ii) and (iii) generally stay weak relative to (i)), so that increasing the rate of ectopic recombination is always disadvantageous, at least under the present assumptions ($R$ not too small).

Overall, TEs thus tend to favor lower recombination rates in the deterministic model, both due to direct cost of ectopic recombination and to the fact that recombination decreases the efficiency of selection against TEs by breaking the positive LD generated by transposition. Increased recombination can be favored in some cases (high $R$, low $\overline{n}$, no correlation between meiotic recombination rates and ectopic recombination) due to the fact that breaking positive LD increases the mean fitness of offspring in the short term, but selection for recombination is extremely weak in this case.

**10. Evolution of recombination: Hill-Robertson effect.** In finite populations, the Hill-Robertson effect tends to generate negative LD among TEs, which may select for increased rates of recombination. The expected change in frequency of the modifier is now given by:

$$\langle \Delta p_m \rangle \approx (s_{\mathrm{dir}} + s_{\mathrm{ind,\,det}} + s_{\mathrm{ind,\,HR}})\, p_m q_m \tag{A107}$$

where $s_{\mathrm{dir}}$ corresponds to direct selection against recombination (first term of equation A96) and $s_{\mathrm{ind,\,det}}$ to deterministic indirect selection (detailed in the previous subsection), while $s_{\mathrm{ind,\,HR}}$ represents indirect selection caused by the Hill-Robertson effect. Assuming $\alpha$, $v \ll u$, and assuming that epistasis among TEs is weak relative to the effective strength of selection acting against insertions (that is, assuming $\beta \ll \alpha + \beta\overline{n} \approx u$), the strength of selection for recombination caused by the Hill-Robertson effect can be obtained from Roze's (2021) analysis of selection for recombination caused by interference between deleterious alleles, the strength of selection against heterozygous mutations ($sh$ in Roze, 2021) being replaced by $u$. One obtains:

$$s_{\mathrm{ind,\,HR}} \approx \frac{\delta R}{N_{\mathrm{e}} R}\, \mathcal{E}[\rho_{ij} \times g(\rho_{mi},\, \rho_{mj},\, \rho_{ij})]\, \frac{\overline{n}^2}{4} \tag{A108}$$

where $g(\rho_{mi},\, \rho_{mj},\, \rho_{ij})$ is a function of the scaled recombination rates $\rho_{mi} = r_{mi}/u$, $\rho_{mj} = r_{mj}/u$ and $\rho_{ij} = r_{ij}/u$ (available in the Supplementary Material), and $\mathcal{E}$ corresponds to the average over all possible pairs of insertion sites $i$ and $j$. Equation A108 yields:

$$\begin{aligned} s_{\mathrm{ind,\,HR}} \approx \frac{\delta R\, u^2}{N_{\mathrm{e}} R^3} \Bigg[ & \int_0^{\frac{R}{2u}} \int_0^{\frac{R}{2u}} (x+y)\, g(x, y, x+y)\, dx\, dy \\ & + \int_0^{\frac{R}{2u}} \int_0^{\frac{R}{2u}} |x-y|\, g(x, y, |x-y|)\, dx\, dy \Bigg] \frac{\overline{n}^2}{2}. \end{aligned} \tag{A109}$$

The first double integral in equation A109 corresponds to the overall effect of pairs of TEs located on opposite sides of the modifier locus on the chromosome, and the second to the overall effect of pairs of loci located on the same side of the modifier.

These integrals can be evaluated numerically using *Mathematica* (see Supplementary Material). When $R \gg u$, they can be approximated by the same integrals evaluated between zero and infinity (Roze, 2021), giving:

$$s_{\text{ind, HR}} \approx 1.8 \frac{\delta R\, u^2\, \overline{n}^2}{8 N_{\text{e}} R^3}. \tag{A110}$$

Note that the effective population size $N_{\text{e}}$ is reduced by background selection effects caused by TEs. Assuming that $R$ is not too small, $N_{\text{e}}$ should stay approximately constant along the chromosome, and can be expressed from classical models of background selection (Hudson and Kaplan, 1995; Charlesworth, 1996), yielding:

$$N_{\text{e}} \approx N \exp\left[-\frac{u\, \overline{n}}{R}\right]. \tag{A111}$$

Finally, when $f$ different TE families (with identical characteristics) are present in the genome, the results above extend to:

$$s_{\text{dir}} \approx -\frac{\delta R}{2} \frac{\tilde{\beta}\, \theta\, f\, \overline{n}^2}{2}, \tag{A112}$$

$$s_{\text{ind, det}} \approx -\frac{\delta R}{8R} \mathcal{E}\left[\frac{r_{ij}}{(r_{mi}+u)\,(r_{mij}+2u)\,(r_{ij}+2u)}\right] u^3 f\, \overline{n}, \tag{A113}$$

$$s_{\text{ind, HR}} \approx \frac{\delta R}{N_{\text{e}} R} \mathcal{E}[\rho_{ij} \times g(\rho_{mi},\, \rho_{mj},\, \rho_{ij})]\, \frac{(f\, \overline{n})^2}{4} \approx 1.8 \frac{\delta R\, u^2 f^2 \overline{n}^2}{8 N_{\text{e}} R^3}, \tag{A114}$$

$$N_{\text{e}} \approx N \exp\left[-\frac{u\, f\, \overline{n}}{R}\right]. \tag{A115}$$

**11. Simulation programs.** The C++ simulation programs (available from Zenodo, doi.org/10.5281/zenodo.7233938) are similar to the programs used in Roze (2021): individuals carry two copies of a linear chromosome with an effectively infinite number of insertion sites. A chromosome is represented by a C++ vector, holding the positions of TEs present on the chromosome (between 0 and 1). When $f$ TE families are segregating, a chromosome is represented by $f$ vectors, holding the positions of TEs from the different families. The initial number of elements from each family is drawn from a Poisson distribution with parameter $n_{\text{init}} = 10$ for each individual, and the position of each TE (on a random chromosome) is drawn from a uniform distribution between 0 and 1 ($n_{\text{init}}$ was set to 100 in some simulations with $\overline{n} = 100$, as explained in the figure legends). Every generation, each TE insertion is eliminated with probability

$v$ (excision rate); transposition then occurs, the number of new elements in a given individual being drawn from a binomial distribution with parameters $u$ and $n$ (where $n$ is the number of elements present in the individual before transposition), while the position of each new element (on a random chromosome) is drawn from a uniform distribution between 0 and 1. To form the next generation of individuals, parents are sampled according to their fitness; during meiosis the number of crossovers is drawn from a Poisson distribution with parameter $R$, and the position of each crossover is drawn from a uniform distribution between 0 and 1. The mean and variance in the number of elements per individual are measured at the beginning of generations (before excision), while $\sum_i {p_i}^2$ is estimated from the mean number of shared insertions between pairs of chromosomes sampled at random from the population (over 1000 chromosome pairs). A second program includes a recombination modifier locus affecting the genetic map length of the chromosome $R$. The modifier is located at the mid-point of the chromosome and mutates at a rate $\mu$ per generation. When a mutation occurs, with probability 0.95 the map length coded by the allele is multiplied by a number drawn from a Gaussian distribution with average 1 and variance 0.04, while with probability 0.05 a number drawn from a uniform distribution between -1 and 1 is added to the value coded by the allele (to allow for large effect mutations), the new value being set to zero if it is negative. The map length $R$ of an individual corresponds to the average between the values coded by its two modifier alleles (additivity). Each simulation lasts $10^6$ generations, $R$ being fixed to 1 during the first 20,000 generations in order to let the number of TEs per individual equilibrate. The mutation rate $\mu$ at the modifier locus was either set to $10^{-3}$ or to $10^{-4}$ depending on simulation runs: when the average chromosome map length $\overline{R}$ at equilibrium is not too small (roughly, $\overline{R} > 0.05$) both values of $\mu$ generally lead to very similar $\overline{R}$, but equilibrium is reached after a larger number of generations when $\mu = 10^{-4}$. However, for small equilibrium values of $\overline{R}$, simulations with $\mu = 10^{-3}$ lead to higher values of $\overline{R}$ than simulations with $\mu = 10^{-4}$, probably due to the fact that mutation is biased towards higher values of $R$ when $R$ is near zero. Therefore, $\mu$ was set to $10^{-3}$ when the predicted evolutionarily stable value of $R$ was higher than 0.05 (in order to reduce execution time), and to $10^{-4}$ when the predicted equilibrium value of $R$ was less than 0.05 (in order to reduce the effect of mutation bias).
